# Pol II degradation activates cell death independently from the loss of transcription

**DOI:** 10.1101/2024.12.09.627542

**Authors:** Nicholas W. Harper, Gavin A. Birdsall, Megan E. Honeywell, Athma A. Pai, Michael J. Lee

## Abstract

Pol II-mediated transcription is essential for eukaryotic life. While loss of transcription is thought to be universally lethal, the associated mechanisms promoting cell death are not yet known. Here, we show that death following loss of Pol II is not caused by dysregulated gene expression. Instead, death occurs in response to the loss of Pol II protein itself, specifically loss of the enzymatic subunit, Rbp1. Loss of Pol II exclusively activates apoptosis, and expression of a transcriptionally inactive version of Rpb1 rescues cell viability. Using functional genomics, we identify a previously uncharacterized mechanism that regulates lethality following loss of Pol II, which we call the **<u>P</u>**ol II **<u>D</u>**egradation-dependent **<u>A</u>**poptotic **<u>R</u>**esponse (**PDAR**). Using the genetic dependencies of PDAR, we identify clinically used drugs that owe their efficacy to a PDAR-dependent mechanism. Our findings unveil a novel apoptotic signaling response that contributes to the efficacy of a wide array of anti-cancer therapies.

## INTRODUCTION

Accurate control of gene expression is critical for all cellular processes. For nuclear protein-coding genes, expression depends on the transcriptional activity of RNA polymerase II (hereafter referred to as Pol II). Pol II activity is considered a life-essential function, and prolonged inhibition of Pol II activity is expected to be universally lethal to cells^1^. In recent years, drugs designed to target the transcriptional machinery have been investigated in clinical settings, especially in the treatment of various cancers^2^. However, despite the importance of Pol II-mediated transcription and clinical interest in drugs targeting the transcriptional cycle, the mechanisms of lethality following transcriptional inhibition remain unclear.

The mechanisms of lethality following Pol II inhibition remain molecularly uncharacterized largely due to the presumption that death in this context is not regulated. Regulated or “active” cell death refers to death induced by active signaling through specific effector enzymes or defined regulatory pathways^3^. In contrast, transcriptional inhibition is generally assumed to cause death due to passive mechanisms; this begins with mRNA decay and the subsequent loss of protein, leading to some unavoidable catastrophic event. Cell death of this type is sometimes referred to as “accidental cell death”, highlighting the use of passive mechanisms, such as decay^4^.

In contrast with the notion that transcriptional inhibition causes an accidental death, recent studies highlight that cells are remarkably capable of buffering against perturbations to the mRNA pool^5–9^. For instance, in normal conditions mRNA concentration scales with cell size; variations in cell size are buffered by tuning Pol II activity or mRNA decay to maintain stable mRNA concentrations^7,10^. Likewise, in perturbed conditions with reduced mRNA production, the effects are also blunted by a reduction in mRNA degradation rates^7,8^. Thus, if cells have the capacity to buffer against the downstream effects of transcriptional inhibition, why do they instead die? To address this question, we aimed to formally characterize the mechanism(s) of lethality following Pol II inhibition in different contexts. We make the unexpected discovery that lethality following Pol II inhibition is not caused by the general decay of mRNA and protein, and instead, is specifically activated by loss of Pol II itself, which initiates a previously uncharacterized apoptotic response.

## RESULTS

### Pol II inhibition exclusively activates an apoptotic cell death

To investigate the mechanisms of cell death by transcriptional inhibition, we focused on triptolide and ⍺-amanitin, two well-validated transcriptional inhibitors that result in the degradation of Pol II^11,12^. We first measured the levels of Rpb1, the largest essential subunit of Pol II. As expected, U2OS cells exposed to a high dose of triptolide rapidly and completely degraded Rpb1, and Rpb1 degradation kinetics could be titrated in a drug dose-dependent manner (Fig. 1a,b and Extended Data FSSig. 1a,b). To validate that Rpb1 protein expression could be used to interpret the transcriptional state, we also measured incorporation of 5-ethynyl uridine (EU) into newly synthesized RNAs, which confirmed the kinetics of transcriptional inhibition following triptolide exposure (Extended Data FSSig. 1c-e). Similar dose and time behavior was also observed following exposure to ⍺-amanitin (Extended Data FSSig. 1f).

**Fig. 1:**
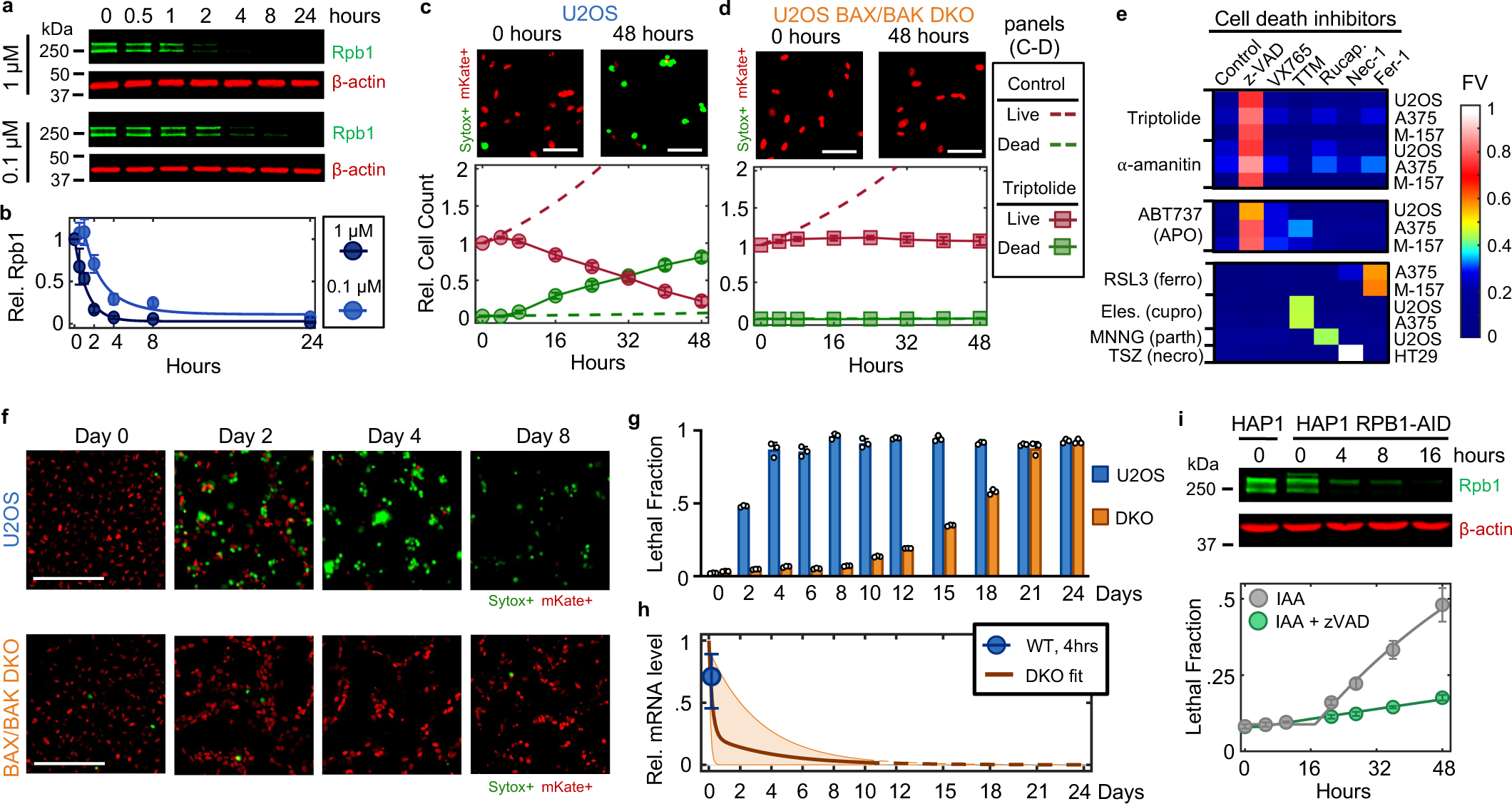
Transcriptional inhibition activates apoptosis prior to dysregulation due to mRNA and protein decay. (**a**) Immunoblot of total Rpb1 protein in U2OS cells following exposure to 1 µM or 0.1 µM Triptolide. Blots are representative of 3 independent biological replicates. (**b**) Quantification of Rpb1 levels shown in panel (a). Mean ± SD are shown. (**c-d**) Live and dead cell kinetics following exposure to 1 µM triptolide for Nuc::mKate2-expressing U2OS cells. Data collected using the STACK assay. mKate+ cells are live, and Sytox+ cells are dead. (c) U2OS. (d) U2OS^BAX-/-/BAK1-/-^ cells (BAX/BAK DKO). Representative images for each genotype are shown above the quantification at the indicated timepoints. Scale bar, 100 µm. Quantified cell numbers depict mean ± SD for 3 independent biological replicates. (**e**) Heatmap of fractional viability (FV) for cells treated with the indicated drugs in the presence/absence of death pathway specific inhibitors. z-VAD-FMK (z-VAD) targets apoptotic caspases; VX765 inhibits the pyroptotic initiator, caspase-1; TTM inhibits cuproptosis (cupro); Rucaparib (Rucap.) inhibits parthanatos (parth); Nec-1 inhibits necroptosis (necro); Fer-1 inhibits ferroptosis (ferro). RSL3, Elesclomol (Eles.), MNNG, and TSZ are canonical activators of the listed death pathways. M-157 = MDA-MB-157. Mean of 3 independent biological replicates is shown. (**f-g**) STACK-based analysis of live and dead cells over extended times following exposure to 1 µM triptolide. (**f**) Representative images from three biological replicates. Scale bar, 275 µm. (**g**) Lethal fraction measured using FLICK following imaging. Data are mean ± SD of 3 biological replicates. (**h**) mRNA levels following exposure to 1 µM triptolide, relative to untreated cells, measured using spike-in normalized RNAseq. Data were normalized to absolute mRNA abundance using polyadenylated ERCC spike-ins. mRNA levels were quantified only for times in which substantial lethality had not yet occurred, which was 4-hours for WT cells, and the first 10 days for DKO cells. Decay kinetics for DKO were fit to a two-term exponential decay function. Range for DKO represents the 50 longest and 50 shortest half-life mRNAs. Dashed lines denote extrapolated expectation from fits. For WT data, median, 25^th^, and 75^th^ percentiles are shown. (**i**) (top) Immunoblot of total Rpb1 protein levels in HAP1 cells, or HAP1-RPB1-AID cells following exposure to 500 µg/mL auxin (IAA, 3-indoleacetic acid). (bottom) Lethal Fraction measurements for HAP1-RPB1-AID cells following 500 µg/mL IAA, in the presence or absence of 50 µM z-VAD. Data collected using the FLICK assay. Data are mean ± SD of 9 independent biological replicates.

To explore cell death in response to Pol II inhibition, we used 1 µM triptolide, which completely inactivated transcription within ∼4 hours (Fig. 1a,b and Extended Data FSSig. 1d). We used the Scalable Time-lapse Analysis of Cell death Kinetics (STACK) assay, which quantifies both live and dead cell populations^13^. We observed that live cells proliferated normally for only the first 4 hours following triptolide exposure, followed by a complete loss of proliferation, coinciding with the initiation of cell death (Fig. 1c). Cell death kinetics at lower doses of triptolide were similar, but with a delay in death onset that matched the later degradation of Pol II (Extended Data FSSig. 2a,b). Similar results were observed with ⍺-amanitin, suggesting that rapid onset of lethality is a general feature of drugs that degrade Pol II (Extended Data FSSig. 2c,d). Taken together, these data reveal that Pol II inhibition activates cell death much more rapidly than previously appreciated, and that cell death is the primary phenotypic outcome of complete transcriptional inhibition.

Since inhibition of Pol II initiated cell death more rapidly than expected, we next aimed to characterize which death mechanism(s) are activated by loss of transcription (Extended Data FSSig. 3a). At least 14 distinct forms of regulated cell death have been identified^14–16^. We reasoned that if cell death following loss of transcription resulted from passive protein loss, rather than an active signaling process, then inhibition of any single death pathway should fail to rescue viability. To test this, we genetically or chemically inhibited each death pathway, one at a time, and observed the effect on cell death following triptolide exposure. To inhibit apoptosis, we knocked out the mitochondrial pore-forming proteins, BAX and BAK (*BAK1*)^17,18^. Remarkably, BAX/BAK double-knockout (DKO) cells were completely resistant to the lethal effects of triptolide (Fig. 1d and Extended Data FSSig. 3b,c). While BAX/BAK DKO cells did not die, these cells were rendered completely non-proliferative within 4 hours of triptolide exposure, suggesting that triptolide had completely inhibited transcription in these cells (Fig. 1d and Extended Data FSSig. 3d,e). Triptolide-induced death was also suppressed using the caspase inhibitor, z-VAD, whereas selective inhibitors of other death pathways did not alter the lethality of triptolide (Fig. 1e). Similar effects were observed following ⍺-amanitin treatment in U2OS cells (Fig. 1e and Extended Data FSSig. 3f,g). Furthermore, inhibiting apoptosis suppressed the lethality of transcriptional inhibitors in a panel of genetically unrelated cell lines (Fig. 1e and Extended Data FSSig. 3h and Supplementary Table 1). These data reveal that transcriptional inhibition – even complete and prolonged transcriptional inhibition – exclusively activates a single cell death pathway, the cell-intrinsic apoptotic response.

### Apoptosis is activated prior to passive loss of apoptotic regulatory proteins

Pol II was completely inhibited at the doses we used, and as expected, RNA levels decreased rapidly following loss of Pol II activity (Extended Data FSSig. 4a-c). However, at the time of death initiation, the RNA pool was only modestly decreased, with the total abundance of mRNAs decayed by ∼25% (Extended Data FSSig. 5a,b). Furthermore, at the protein level, we observe no change in the expression of canonical apoptotic regulatory proteins 4 hours after triptolide exposure (Extended Data FSSig. 5c-f). Thus, the rapid initiation of apoptosis following Pol II inhibition could not be easily explained from the passive loss of mRNA or downstream decrease in protein expression.

Cellular behaviors did not appear to be dysregulated, even following a more substantial loss of RNAs. For instance, we observed similar RNA decay rates in U2OS and U2OS-BAX/BAK DKO cells (Extended Data FSSig. 6a). However, 4 days following triptolide exposure, when U2OS were completely dead, BAX/BAK DKO cells display no overt signs of dysregulation, despite having lost more than 90% of the RNA pool (Fig. 1f-h and Extended Data FSSig. 6a-d). Even 4 days after triptolide exposure, BAX/BAK DKO cells: were capable of re-adhering to plates following trypsinization, produced roughly normal levels of ATP, and could actively signal following exposure to external stresses, such as DNA damage (Extended Data FSSig. 7a-g). Normal behaviors were maintained in BAX/BAK DKO cells through ∼10 days of triptolide exposure, after which, cells began dying, with the entire population dying by 24 days (Fig. 1g). Notably, the passive death observed in BAX/BAK DKO cells was coincident with the complete loss of all mRNAs, as evaluated using absolute mRNA quantification (Fig. 1h and Extended Data FSSig. 8a,b). These data reveal that accidental cell death following loss of mRNA does indeed occur, but is temporally distinct from the rapid death activated by Pol II loss in cells that are proficient for activating apoptosis.

One potential explanation for these unexpected results is that the lethality induced by triptolide and ⍺-amanitin stems from an off-target mechanism, unrelated to Pol II inhibition. To address this hypothesis, we evaluated the effects Pol II degradation in HAP1 cells, in which the endogenous RPB1 gene (*POL2RA*) is tagged with an auxin-inducible degron (HAP1-RPB1-AID) (Fig. 1i, top)^19^. While auxin had no effect on wild-type HAP1 cells, HAP1-RPB1-AID cells displayed rapid cell death that initiated shortly after Pol II degradation (Fig. 1i and Extended Data FSSig. 9a). Furthermore, the lethality of Rpb1 degradation could be potently suppressed by co-treatment with z-VAD, consistent with the results observed for triptolide and ⍺-amanitin (Fig. 1i). Taken together, these data reveal that apoptosis following Pol II inhibition occurs in response to an active process that is distinct from the effects of global loss of mRNA.

### Loss of mRNA production is not correlated with the induction of apoptosis

While apoptotic-proficient cells die before experiencing global loss of mRNA, these cells still experience a complete loss of extremely short-lived transcripts (Fig. 1h, and Extended Data FSSig. 6b). Mcl-1 is a potent negative regulator of apoptosis, that is unique among apoptotic regulators in that it is rapidly turned over at both the mRNA and protein levels^20–23^. Indeed, Mcl-1 is rapidly lost following transcriptional inhibition at both the mRNA and protein levels (Extended Data FSSig. 10a,b). However, U2OS cells were insensitive to chemical inhibition or genetic knockout of *MCL1*, and loss of *MCL1* did not alter sensitivity to triptolide (Extended Data FSSig. 10c,d).

We next sought to more generally test if loss of short-lived RNAs following transcriptional inhibition causes apoptosis. One simple prediction of this model is that drugs that inhibit Pol II activity faster will initiate cell death proportionally faster. Thus, coordinated variation in drug-induced decay and the subsequent activation of cell death may help to identify causal events. To capture variation in Pol II inhibition kinetics, we profiled transcriptional inhibitors that function with different inhibition timing: flavopiridol, THZ1, triptolide, actinomycin-D, α-amanitin, and ethynylcytidine (Extended Data FSSig. 11a-d)^24^. To quantify the kinetics of Pol II inhibition for these drugs, we used immunoblotting of Rpb1 to distinguish between the levels of active and inactive Pol II. Rpb1 migrates as two distinct bands. The upper band, called Pol II-o, migrates slowly due to hyper-phosphorylation of the highly conserved series of heptapeptide repeats in the regulatory carboxyl-terminal domain (CTD)^25^. Importantly, Pol II-o is the actively elongating pool of Pol II^26^. The lower band, called Pol II-a, represents all other inactive forms of Pol II, including preinitiation complex (PIC) bound, promoter paused, early pause released, and free Pol II^27^. We observed significant variation in the kinetics of Pol II inhibition, with some drugs, such as the Cdk9 inhibitor flavopiridol, completely inhibiting Pol II activity within minutes, while other indirect inhibitors required several hours to completely inactivate Pol II (Fig. 2a-b and Extended Data FSSig. 11a-d). However, the fastest-acting transcriptional inhibitors were not particularly fast at activating cell death. The two fastest transcriptional inhibitors, flavopiridol and THZ1, were, in fact, the two slowest activators of cell death (Fig. 2b). Overall, we found no correlation between the timing of Pol II inhibition and timing of cell death (Fig. 2c,d and Supplementary Table S2). These data suggest that the death induced by Pol II inhibition does not result from the loss of short-lived mRNAs.

**Fig. 2:**
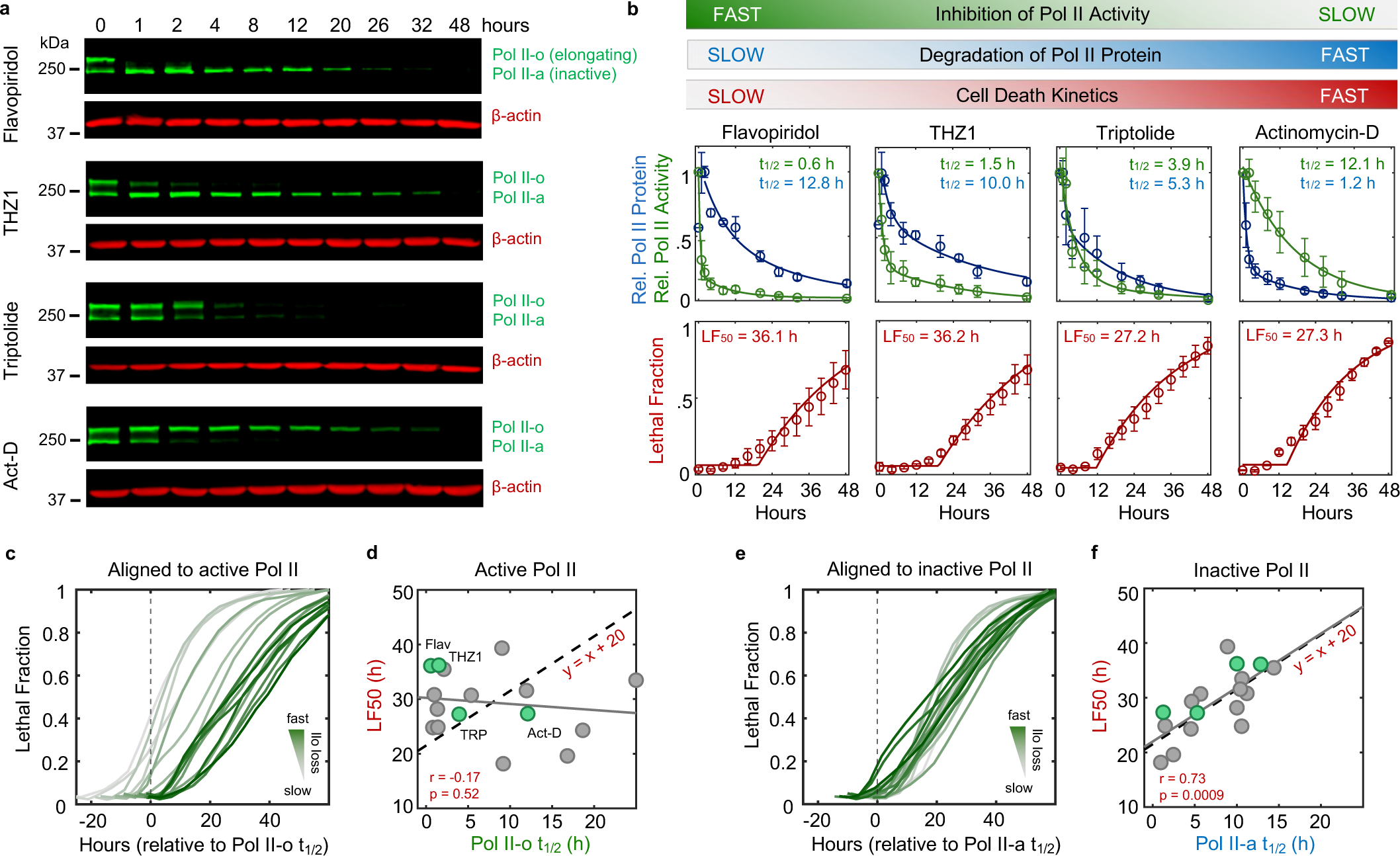
Loss of Pol II protein – but not loss of Pol II activity – correlates with onset of apoptotic death. (**a**) Immunoblots of total Rpb1 protein following exposure to either flavopiridol (10 µM), THZ1 (1 µM), triptolide (0.1 µM), or actinomycin-D (Act-D, 0.1 µM). Blots are representative of three independent biological replicates. (**b**) (Top panel) Quantification of Pol II-o (elongating) and Pol II-a (inactive) decay kinetics from immunoblots described in (a). Data fit to a two-term exponential decay equation. t_1/2_ denotes time to 50% decay. (Bottom panel) Cell death kinetics quantified using the STACK assay. LF_50_ denotes time to 50% max observed death. For both panels, data are mean ± SD, n = 3 independent biological replicates. (**c-d**) Relationship between loss of Pol II activity and activation of cell death. **(c)** Cell death kinetics for 17 unique transcriptional inhibitor drug-dose combinations, aligned to t_1/2_ for loss of active Pol II (dashed vertical line). Cell death was quantified using the STACK assay, and the mean of 3 independent biological replicates measured every four hours is shown. (**d**) For data in (c), correlation between t_1/2_ for loss of active Pol II and time to 50% lethality (LF50). Solid gray line denotes fit to a linear regression model. Black dashed line denotes a direct x = y relationship, shifted by a constant to account for a lag time between drug activity and cell death. **(e-f)** Relationship between loss of inactive Pol II and activation of cell death. **(e)** Same data as (c), except aligned to t_1/2_ for degradation of inactive Pol II (dashed vertical line). **(f)** For data in (e), correlation between t_1/2_ for degradation of inactive Pol II and time to 50% lethality (LF50).

The effects of transcriptional inhibitors could be phenocopied by degrading Rpb1 using an auxin-inducible degron (Fig. 1i). Thus, we considered other mechanisms by which loss of Pol II could activate apoptotic death. While previous studies have often focused on how transcriptional inhibitors target actively elongating Pol II-o, our data highlight an underappreciated ability of these drugs to modulate the inactive Pol II-a pool as well (Fig. 2a). Notably, while loss of Pol II *activity* was not correlated with activation of death, we observed a strong positive correlation between timing of cell death and timing of degradation for the *inactive* Pol II protein (Fig. 2e,f and Extended Data FSSig. 11e). Furthermore, the relationship between inactive Pol II decay and cell death was directly proportional, such that loss of inactive Pol II consistently preceded cell death by a fixed amount of time (Fig. 2f). While only a correlative result, these data propose an alternative model whereby death following transcriptional inhibition is not related to the inhibition of Pol II activity and the subsequent loss of mRNA, but instead may result directly from the degradation of Pol II protein.

### Loss of Pol II protein, not loss of mRNA, activates apoptosis

A critical challenge for determining causality in this context is the mutual dependence between Pol II protein and its enzymatic function: degrading Pol II invariably inhibits Pol II activity, and the existence of Pol II protein relies on Pol II-mediated transcription. Indeed, we were unable to identify any compound capable of completely inhibiting transcription without also inevitably degrading Pol II protein. To overcome this challenge, we developed three complementary strategies for uncoupling the inhibition of Pol II activity and the degradation of Pol II protein (Fig. 3a, i-iii). First, we acutely degraded Pol II without inducing subsequent loss of mRNAs, by exposing cells to triptolide while concurrently inhibiting key mediators of RNA degradation (Fig. 3a-i). NUP93 and EXOSC5 have been shown to be critical mediators of mRNA concentration homeostasis, and acute knockdown of these proteins leads to an accumulation of mRNA^7^. As expected, the mRNA pool in cells lacking NUP93 was stabilized, and significantly more resistant to triptolide-induced mRNA loss than control cells (Fig. 3b). Despite the stabilized mRNA pool, however, triptolide activated cell death to similar levels – and with similar kinetics – in control and NUP93-KO cells (Fig. 3c). Similar results were observed in EXOSC5-KO cells (Extended Data FSSig. 12a,b). These data again suggest that cell death following Pol II inhibition cannot be explained by the passive loss of mRNA alone.

**Fig. 3:**
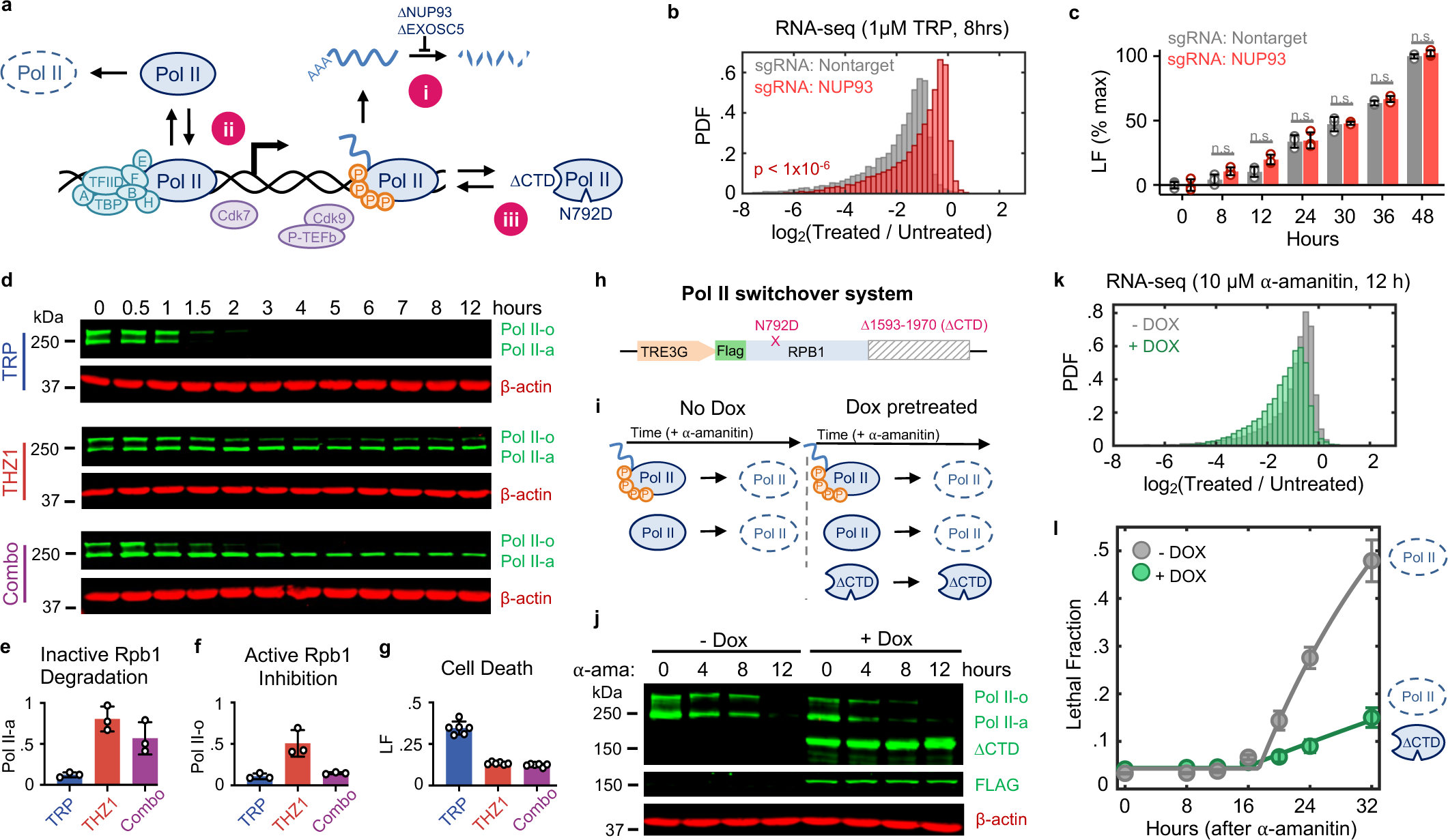
Exogenous expression of inactive Pol II rescues viability following Pol II degradation. (**a**) Schematic describing three approaches (i – iii) for uncoupling inhibition of Pol II enzymatic activity from degradation of Pol II protein. (**b**) Histogram of spike-in normalized absolute mRNA fold changes following 8-hr exposure to 1 µM triptolide (TRP) in U2OS cells expressing sgRNA targeting NUP93 or nontargeting sgRNA. Two-sided KS test *p* value is shown. (**c**) Comparison of lethal fraction (LF) levels between cell types in (b) at various time points following exposure to 1 µM TRP, measured using the FLICK assay. Data normalized to max LF in cells expressing nontargeting sgRNA. Data are mean ± SD for n = 3 independent biological replicates. Wilcoxon rank sum p value shown (n.s. > 0.05). (**d**) Immunoblot of total Rpb1 levels in U2OS cells following exposure to 1 µM TRP (top), 0.316 µM THZ1 (middle), or the combination of the two (bottom). Blots shown are representative of three independent biological replicates. (**e-f**) Quantification of Rpb1 levels after 4-hr drug exposure (associated with panel d). (**e**) inactive (ll-a). (**f**) active (ll-o). **(g)** LF at 24-hrs, measured using the FLICK assay in identical conditions. Data are mean ± SD for n = 6 independent biological replicates. (**h-l**) Pol II switchover system used to evaluate function of inactive Pol II. **(h)** Diagram of a Pol II construct featuring a doxycycline-inducible promoter, FLAG tag, α-amanitin resistance mutation (N792D), and deletion of amino acids 1593-1970 (ΔCTD). **(i)** Experimental logic behind the Pol II switchover system. Tet-ON U2OS cells expressing the N792D/ΔCTD construct are exposed to 10 µM ⍺-amanitin in the presence or absence of 2 µg/mL doxycycline (Dox). Dox added 48-hrs prior to ⍺-amanitin to allow for expression of N792D/ΔCTD prior to degradation of the endogenous Pol II-o/a. (**j**) Immunoblots of Rpb1 following exposure to 10 µM ⍺-amanitin. (**k**) Histogram of spike-in normalized absolute mRNA fold changes following 12-hr exposure to 10 µM ⍺-amanitin. (**l**) LF kinetics following 10 µM ⍺-amanitin. LF measured using FLICK. Mean ± SD shown, n = 5 independent biological replicates.

Second, we sought to buffer against triptolide-induced degradation of Pol II, while maintaining transcriptional inhibition (Fig. 3a-ii). Triptolide inhibits XPB, the helicase subunit of TFIIH^11^. Triptolide-induced degradation of Pol II has been shown to require Cdk7 activity, presumably because Cdk7 phosphorylation of Pol II facilitates removal of the pre-initiation complex, resulting in the release of Pol II from chromatin, enabling Pol II degradation^27^. Consistent with this, co-treatment with the Cdk7 inhibitor, THZ1, blocks Pol II degradation following high-dose triptolide exposure (Fig. 3d,e). Importantly, THZ1 is itself a transcriptional inhibitor, and the combination of triptolide and THZ1 was as good at inhibiting Pol II activity as either drug alone (Fig. 3d,f). Despite the potently inhibited transcriptional activity, we find that the addition of THZ1 suppresses the lethality of triptolide while maintaining potent transcriptional inhibition (Fig. 3d,g). This paradox – in which the lethal effects of transcriptional inhibition can be suppressed by adding another transcriptional inhibitor – is in line with a model whereby lethality results from the degradation of Pol II protein.

Finally, we asked if exogenous expression of a functionally inactive Pol II mutant protein was sufficient to rescue viability following degradation of the endogenous Pol II pool (Fig. 3a-iii). To accomplish this, we designed a Pol II “switchover” system (Fig. 3h). This system features a doxycycline-inducible *RPB1* transgene completely devoid of the regulatory carboxy-terminal domain (CTD) that is required for productive transcription^28–31^. In addition, the *RBP1* fragment contains a known ⍺-amanitin resistance mutation, N792D, allowing the selective degradation of endogenous Pol II^12^. Conceptually, this system allows us to directly compare cell death outcomes following transcriptional inhibition, using isogenic cells that either have, or do not have, a pool of non-functional Pol II protein (Fig. 3i). To test this system, we stably expressed the doxycycline-inducible *RPB1*-N792D-ΔCTD construct in U2OS cells expressing a Tet-On trans-activation protein. These cells were pretreated with- or without doxycycline for two days to enable expression of Rpb1-N792D-ΔCTD, followed by acute degradation of endogenous Pol II using 10 µM ⍺-amanitin. As expected, complete loss of endogenous Pol II was observed following ⍺-amanitin exposure, in both doxycycline-pretreated and untreated conditions (Fig. 3j). However, only cells pretreated with doxycycline expressed a truncated Rpb1 protein that did not respond to ⍺-amanitin (Fig. 3j). Expression of Rpb1-N792D-ΔCTD did not inhibit the loss of mRNA following ⍺-amanitin treatment, validating that the CTD is required for productive transcription (Fig. 3k, and Extended Data FSSig. 13a). Importantly, expression of Rpb1-N792D-ΔCTD potently suppressed ⍺-amanitin-induced cell death (Fig. 3l). Expression of Rpb1-N792D-ΔCTD had no effect on the lethality of unrelated apoptotic agents, and rendering the Rpb1-ΔCTD protein sensitive to ⍺-amanitin abolished its ability to suppresses cell death (Extended Data FSSig. 13b,c). These experiments demonstrate that cell death following transcriptional inhibition is caused by the degradation of Pol II protein, and not loss of Pol II activity.

### Apoptosis following Pol II degradation requires PTBP1 and BCL2L12

We next aimed to characterize how decreased Pol II levels are sensed and communicated to initiate apoptosis. Considering the lack of prior knowledge for which processes respond to loss of Pol II protein, we evaluated this question using an unbiased genome-wide screening approach. We used Cas9-expressing U2OS cells, and the TKOv3 sgRNA library, to evaluate the effect of all single gene knockouts on triptolide-induced lethality^32,33^. To identify genes that specifically regulate cell death, rather than proliferation, we developed a simple experimental approach to mechanically separate live cells from apoptotic corpses, such that both populations could be sequenced independently (Fig. 4a, Extended Data FSSig. 14a-f and Supplementary Table 3).

**Fig. 4:**
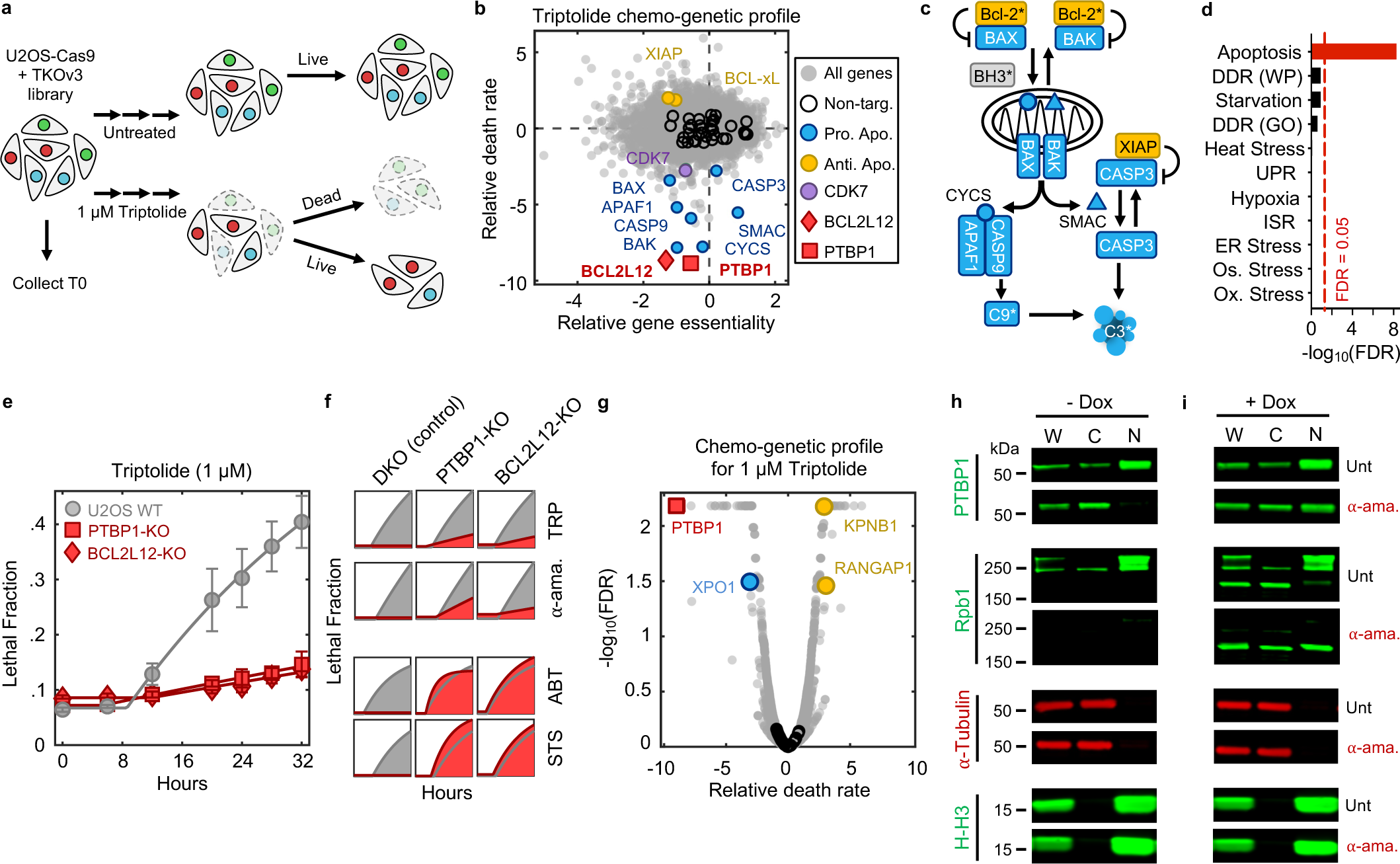
Apoptosis following Pol II degradation exhibits unique genetic dependencies. (**a**) Schematic of chemo-genetic profiling to identify cell death regulatory genes. (**b**) Gene-level chemo-genetic profiling data for U2OS cells treated with 1 µM triptolide. Key death regulatory genes and non-targeting controls are highlighted. (**c**) Simplified schematic of the cell intrinsic apoptotic pathway. Bcl-2* denotes the general anti-apoptotic Bcl-2 family members, whereas BH3* denotes the pro-apoptotic BH3-only family members. Regulators in blue/yellow were identified in the chemo-genetic profiling data. (**d**) Enrichment of various stress response pathways in genes whose knockout significantly suppresses triptolide-induced cell death, as identified in the chemo-genetic profiling. Significance was determined using a one-tailed Fisher’s exact test corrected for multiple comparisons using the Benjamini-Hochberg (BH) FDR. (**e**) Cell death kinetics measured using FLICK following 1µM triptolide in U2OS wild-type cells, PTBP1-KO cells (red, square), BCL2L12-KO cells (red, diamond). Data are mean ± SD for n = 6 independent biological replicates. (**f**) PTBP1 and BCL2L12 dependence for drugs that degrade Pol II, or canonical apoptotic agents: triptolide (TRP): 1 µM; ⍺-amanitin (⍺-ama.): 10 µM; ABT-199 (ABT): 100 µM; Staurosporine (STS): 0.5 µM. Drugs tested in U2OS-PTBP1-KO, U2OS-BCL2L12-KO, or U2OS BAX/BAK double knockout (DKO). Lethal fraction kinetics shown with area under the kinetic curve colored: grey = wild-type cells, red = DKO, PTBP1-KO, or BCL2L12-KO cells, as indicated. Areas are based on the mean of 6 independent biological replicates. **(g**) Drug-induced death rates from chemo-genetic profiling of triptolide, with genes highlighted that regulate PTBP1 translocation: nuclear import factors in yellow; nuclear export factors in blue. Non-targeting genes in black. (**h-i**) Nuclear/cytoplasmic fractionation to assess PTBP1 localization following exposure to ⍺-ama. W = whole cell extract; C = cytoplasmic extract; N = nuclear extract. Immunoblots for Rbp1 shown to validate ⍺-ama. function; ⍺-Tubulin and Histone-H3 (H-H3) shown as markers of cytoplasmic and nuclear proteins, respectively. **(h)** Data are from U2OS-RBP1-N792D/ΔCTD cells exposed to 10 µM ⍺-amanitin for 12 hours. **(i)** As in panel (h), except samples collected from U2OS-RBP1-N792D/ΔCTD cells exposed to doxycycline (Dox) for 48 hours prior to exposure to ⍺-amanitin.

Our screen identified more than 100 genes whose knockout significantly reduced the level of triptolide-induced cell death (Fig. 4b and Supplementary Table 4). Remarkably, this cohort of genes included every gene encoding an essential pro-apoptotic regulatory protein (Fig. 4b,c). We also identified key negative regulators of apoptosis, such as *XIAP* and BCL-xL (*BCL2L1*), which increase triptolide-induced lethality when deleted (Fig. 4b,c). Our screen also identified triptolide-specific genetic dependencies, including *CDK7*, in line with our finding that the Cdk7 inhibitor, THZ1, also rescues triptolide-induced cell death (Fig. 4b and 3g).

To clarify the mechanism by which Pol II degradation activates apoptosis, we began by exploring if the regulators of triptolide-induced cell death were enriched for genes involved in known stress response pathways. As expected, given that Pol II degradation exclusively activates apoptosis, the hits from our screen were strongly enriched for apoptotic regulatory genes (Fig. 4d); however, we did not identify any other known stress response pathways that might facilitate activation of apoptosis (Extended Data FSSig. 15a). Due to the proximity of Pol II to DNA, we paid special attention to regulators of the DNA damage response (DDR). However, the DDR did not appear to be involved in triptolide-induced cell death: no regulators of the DDR were hits in our screen, triptolide failed to activate DDR signaling, and knocking out TP53, an essential regulator of DDR-induced apoptosis^34^, had no effect on triptolide-induced cell death (Extended Data FSSig. 15b-d).

We next explored if Pol II degradation activates apoptosis through a unique and yet undescribed process. Surprisingly, the two genes whose deletion most strongly suppressed triptolide-induced cell death were not core apoptotic effectors, but PTBP1 and BCL2L12 (Fig. 4b and Extended Data FSSig. 16a). BCL2L12 is a poorly understood protein with an undefined role in apoptotic activation, even though the Bcl2 family has been extensively studied in the context of apoptosis^35^. PTBP1 is a multi-functional RNA binding protein, with well-established roles in alternative splicing and activation of internal ribosome entry segments (IRESs)^36–38^. To validate the role of these genes in triptolide-induced cell death, we generated *PTBP1* and *BCL2L12* knockout cells in the U2OS genetic background. Knocking out either *PTBP1* or *BCL2L12* suppressed triptolide-induced cell death, validating the results of our screen (Fig. 4e).

Notably, our screen identified every essential apoptotic effector downstream of mitochondrial outer membrane permeabilization; however, *BCL2L12* was the only Bcl-2 homology (BH) domain-containing gene that suppressed death (Extended Data FSSig. 16b). These data suggest that PTBP1 and BCL2L12 may play a pro-apoptotic role, specifically in the context of Pol II degradation. To test this idea, we explored the effect of *PTBP1* or *BCL2L12* knockout in other drug contexts. As we observed for triptolide, knocking out *PTBP1* or *BCL2L12* suppressed lethality following exposure to other Pol II inhibitors, such as α-amanitin (Fig. 4f and Extended Data FSSig. 16c,d). In contrast, knocking out *PTBP1* or *BCL2L12* failed to suppress lethality in the context of canonical apoptotic agents, such as the BH3 mimetic, ABT-199, or Staurosporine (Fig. 4f and Extended Data FSSig. 16e,f). As expected, all apoptotic drugs failed to activate death in BAX/BAK DKO cells (Fig. 4f and Extended Data FSSig. 16c-f). Together, these data reveal that apoptotic death induced by Pol II degradation requires PTBP1 and BCL2L12.

### Nuclear-to-cytoplasmic translocation of PTBP1 regulates death

Knocking out *PTBP1* did not alter Pol II degradation, nor the triptolide-induced loss of transcriptional activity, or the decay of mRNAs following triptolide exposure (Extended Data FSSig. 17a-c). These data rule out the trivial explanation that loss of *PTBP1* inhibits triptolide activity. Furthermore, triptolide-induced changes in gene expression in U2OS and PTBP1-KO cells were nearly indistinguishable, and the shared expression changes between U2OS and PTBP1-KO cells included apoptotic regulatory mRNAs, such as *MCL1*, which were also rapidly lost at the protein level in both cell types (Extended Data FSSig. 17d-f).

Thus, we next explored the mechanisms by which PTBP1 could regulate apoptosis following Pol II degradation. PTBP1 is an RNA binding protein. Prior studies have reported a post-transcriptional function for PTBP1, which can promote TNF-induced apoptosis by activating IRESs to facilitate translation of apoptotic effector proteins, such as APAF1^39,40^. However, we observed that APAF1 protein expression was not altered during triptolide exposure, and APAF1 was expressed to similar levels in WT and PTBP1-KO cells (Extended Data FSSig. 17f). The best described functions for PTBP1 are in regulating alternative mRNA splicing^41–44^. Intriguingly, the influence of PTBP1 on triptolide-induced lethality also does not appear to be related to its role in splicing. Other regulators of alternative splicing did not rescue the lethality of triptolide (Extended Data FSSig. 18a). Additionally, although we observed alternative splicing of known PTBP1-dependent splicing events in PTBP1-KO cells, triptolide did not affect PTBP1-dependent splicing, and splicing changes induced by triptolide were unaffected in PTBP1-KO cells (Extended Data FSSig. 18b-f)^36^.

Many RNA binding proteins, such as PTBP1, are localized to the nucleus, and accumulate in the cytoplasm following transcriptional inhibition, though the precise reason for this is unclear^45,46^. Curiously, triptolide-induced lethality was rescued by knocking out the nuclear exporter *XPO1*; and conversely, knocking out nuclear import factors, *RANGAP1* or *KPNB1*, sensitized cells to triptolide-induced lethality (Fig. 4g). Furthermore, we observed PTBP1 translocation to the cytoplasm shortly following triptolide or α-amanitin-induced Pol II degradation (Fig. 4h and Extended Data FSSig. 19a,b). Finally, expression of inactive Pol II (Rpb1-N792D-DCTD) blocked the cytoplasmic translocation of PTBP1 and suppressed the lethality of α-amanitin (Fig. 4i and Extended Data FSSig. 19c). These data suggest that the cytoplasmic trafficking of PTBP1 is important for its death regulatory function.

We performed RNAseq on cytoplasmic and nuclear fractions separately in WT and PTBP1-KO cells, and found no evidence that the altered localization of PTBP1 is associated with a change in mRNA localization that can causally explain triptolide-induced cell death (Extended Data FSSig. 19d-f). A cross-comparison of our RNAseq and chemo-genetic profiling datasets further revealed no potential PTBP1-dependent transcriptional effects that could causally explain triptolide-induced cell death (Extended Data FSSig. 19g,h). These data further support our general finding that loss of mRNA cannot causally explain cell death following transcriptional inhibition. Moreover, these data support the notion that PTBP1 is a genuine regulator of apoptosis, specifically in the context of Pol II degradation, and that its mechanism in this context is likely unrelated to its currently known molecular functions.

### Multiple anti-cancer drugs owe their lethality to Pol II degradation

We next aimed to investigate if activation of a Pol II degradation-dependent cell death could be valuable in a therapeutic context. To address this question, we focused on clinically relevant compounds – including those not conventionally considered transcriptional inhibitors – and determined the degree to which the lethality of these drugs depended on genes required for triptolide-induced cell death. We generated a simple “Transcriptional Inhibition Similarity” (TIS) score to quantitatively evaluate the functional similarity of a given drug to the canonical Pol II degrader, triptolide (Fig. 5a,b). We applied this signature-based strategy across 46 diverse compounds, covering 16 clinically relevant drug classes (Fig. 5c and Extended Data FSSig. 20a-i). Strikingly, a wide array of TIS scores were observed across the dataset, with variation both across and within drug classes (Fig. 5c and Extended Data FSSig. 21 and Supplementary Table 5). High TIS scores were observed for most, but not all, drugs that directly target Pol II or transcriptional Cdk proteins (Fig. 5c). Additionally, drugs that induce proteotoxic stress, such as Brefeldin A, Thapsigargin, or Tunicamycin, were quite lethal, but featured low TIS scores, further confirming that Pol II degradation activates death in a manner that is distinct from general dysregulation of protein levels (Extended Data FSSig. 21). Collectively, these data highlight that clinically relevant drugs covering disparate annotated mechanisms share the genetic dependencies observed for transcriptional inhibitors.

**Fig. 5:**
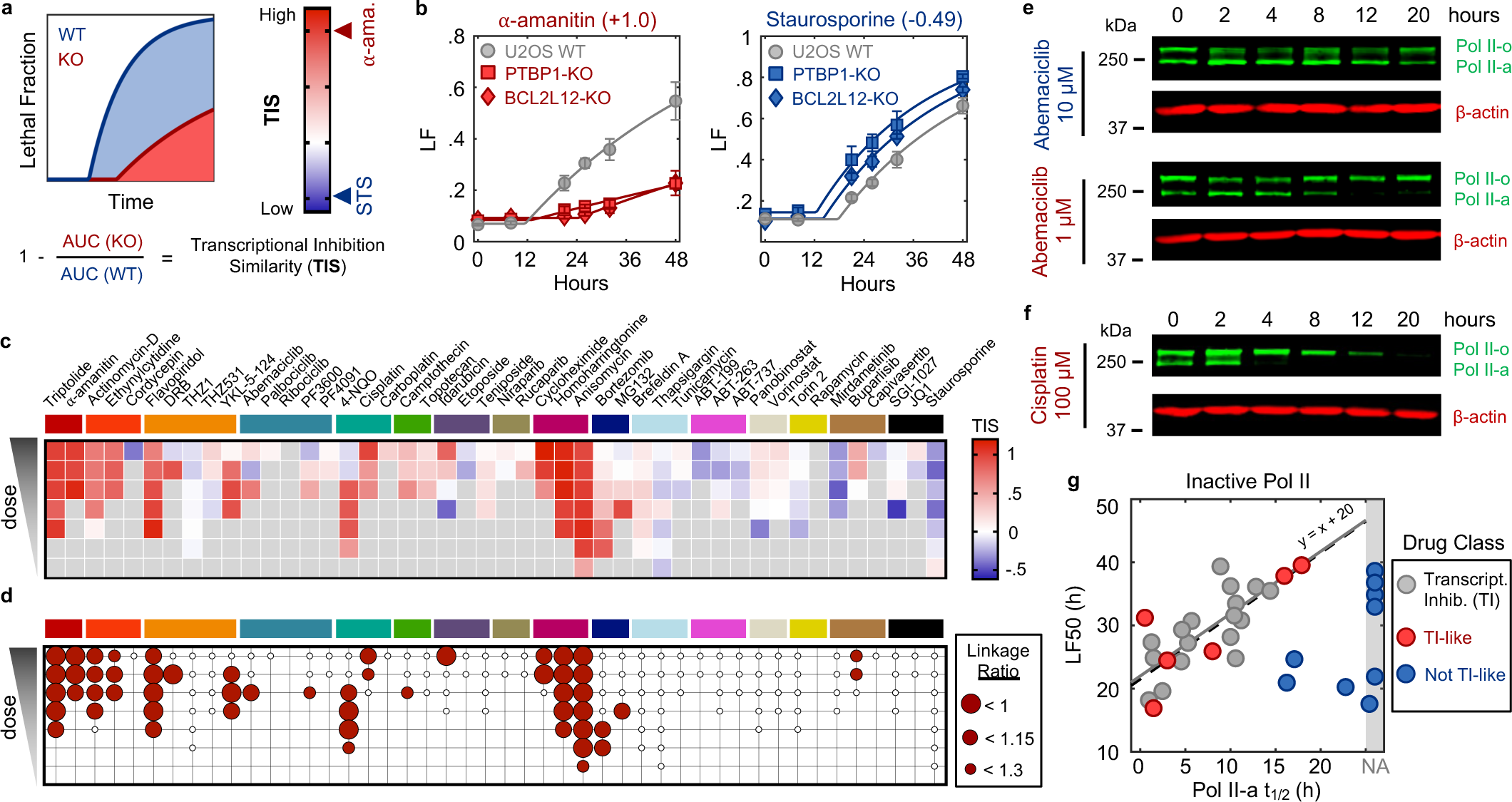
Commonly used anti-cancer drugs owe their lethality to Pol II degradation. (**a**) Schematic illustrating the strategy to quantify the degree to which the lethality of a given drug is dependent on the genetic dependencies observed for Pol II degraders (*i.e.*, PTBP1 and BCL2L12). (**b**) Cell death kinetics in U2OS cells for a high scoring drug (10 µM ⍺-amanitin, TIS = 1.0) and a low scoring drug (3.16 µM Staurosporine, TIS = -0.49). Data collected using FLICK. Data are mean ± SD for n = 3 independent biological replicates. (**c**) Heatmap depicting the Transcription Inhibition Similarity (TIS) score across a 7-point half-log dose range for 46 compounds encompassing diverse established mechanisms of action and clinical utility. Data collected in U2OS cells. Grey boxes are for non-lethal doses of each drug. Compounds are ordered and colored by drug class. See methods for exact doses tested and drug class definitions. (**d**) Binary classifier, defining each drug-dose condition as “Transcriptional Inhibition-like” (TI-like) or not. Classifier results are ordered as in (c). Red circles denote TI-like mechanism of lethality. (**e**) Immunoblot of total Rpb1 levels in U2OS cells following exposure to high dose Abemaciclib (classified as low TIS), or a ten-fold lower dose (classified as high TIS and TI-like mechanism of lethality). (**f**) Immunoblot of total Rpb1 levels in U2OS cells following exposure to high dose Cisplatin. (**g**) Relationship between timing of cell death and timing of Pol II-a loss. Data in gray denote established transcriptional inhibitors, as shown in Fig. 2f. Red and blue data points represent tested TI-like compounds and not TI-like compounds, respectively. Black dashed line denotes a direct x = y relationship, shifted by a constant to account for a lag time between drug activity and cell death.

Because each drug was evaluated across a range of doses, our data revealed an unexpectedly large degree of variation in the TIS score across doses of a drug. To more precisely identify drug-dose pairs that owe their lethality to a triptolide-like mechanism, we developed a binary classifier to evaluate our TIS data. We used a conservative probabilistic nearest neighbors-based classification approach, in which a drug-dose pair was classified as “Transcriptional inhibition-like” only if its proximity to validated Pol II degrading drugs is within the neighborhood occupied by the Pol II degraders, triptolide and α-amanitin (Extended Data FSSig. 22a)^47,48^. Our classifier identified significant variation in drug mechanisms of lethality, particularly across drugs that induce DNA damage (Fig. 5d). For instance, Idarubicin, a Topoisomerase II inhibitor, was lethal at the highest 4 tested doses, but was only “TI-like” at the highest dose (Fig. 5d and Extended Data FSSig. 21). Conversely, the UV radiation-mimetic, 4-NQO, was lethal at nearly all tested doses, but strikingly, was only TI-like at intermediate doses (Fig. 5d and Extended Data FSSig. 21).

To validate our classifier, we next tested whether drugs classified as TI-like also degraded Pol II, and if this behavior was restricted to the appropriate doses. We focused on Abemaciclib, given the unexpected dose dependence, in which our classifier predicts Pol II degradation at only a single intermediate dose, and paradoxically, not at high doses (Extended Data FSSig. 22b). Consistent with our classifier, we found that Pol II is degraded following exposure to 1 µM Abemaciclib, but not when exposed to a ten-fold higher dose (Fig. 5e). These distinctions are particularly surprising given that Abemaciclib is lethal at both doses (Supplementary Table 5). Similar paradoxical dose-dependent Pol II degradation was observed for other drugs, including 4NQO, the Cdk2/4/6 inhibitor PF-06873600 (PF3600), and Idarubicin, (Extended Data FSSig. 22c-i).

Due to its widespread clinical use, we next focused on Cisplatin, a DNA cross-linking drug used in the treatment of bladder, head and neck, lung, ovarian, and testicular cancers, among others^49^. Strikingly, our classifier predicts that Cisplatin has a TI-like mechanism of lethality at all doses that activate high levels of cell death (Fig. 5d and Extended Data FSSig. 23a). This was surprising, given that DNA damage can independently activate apoptosis through the DDR, which is observed for other drugs in this dataset (Extended Data FSSig. 21 and 23a). Consistent with our classifier, we observe Pol II degradation following Cisplatin exposure at all doses that induce substantial lethality (Fig. 5f and Extended Data FSSig. 23b-d). These data suggest that the lethality of several DNA damaging agents may hinge upon a Pol II degradation-dependent apoptotic response in both a drug- and dose-dependent manner.

Finally, to further validate that Pol II degradation is responsible for the lethality of TI-like compounds, we explored other features that are unique to this mechanism. In addition to the dependence on PTBP1, BCL2L12, and the degradation of inactive Pol II protein, death kinetics represent another critical feature of this mechanism. We demonstrated that canonical transcriptional inhibitors activate death with consistent timing following loss of the inactive form of Pol II (Fig. 2f). For drugs classified as “TI-like”, cell death kinetics were well-correlated with the kinetics of inactive Pol II degradation (Fig. 5g). Cell death kinetics for negative control drugs showed no relationship, and no correlation was observed between cell death timing and loss of Pol II activity (Fig. 5g and Supplementary Table 6). These data reveal that the mechanisms of drug-induced lethality vary in unexpected and non-trivial ways across dose, and more specifically, that Pol II degradation is a common mechanism of lethality across many unrelated drug classes.

## DISCUSSION

The fact that transcriptional inhibition is lethal to cells is hardly surprising. However, the common assumption to date has been that prolonged loss of transcription is toxic to cells due to global loss of mRNA and protein. We show that death following transcriptional inhibition initiates much more rapidly than previously appreciated, well before cells are experiencing stress from the loss of mRNA or protein. Furthermore, we demonstrate that death occurs in response to an active signaling process, which unexpectedly, is initiated by loss of the Pol II protein, not the loss of Pol II transcriptional activity, nor the general decay in mRNA or protein levels. Therefore, we propose that this previously uncharacterized process be termed the **<u>P</u>**ol II **<u>D</u>**egradation-dependent **<u>A</u>**poptotic **<u>R</u>**esponse (**PDAR**).

Why has this phenotype not been observed previously? Notably, the central issue is simply related to analytical resolution. Specifically, while apoptotic-deficient cells remain 100% viable following Pol II degradation, these cells also cannot proliferate. Common drug response metrics misinterpret the level of lethality in this context, because a non-proliferative drug-treated population is already very small when compared to a rapidly dividing untreated population (Extended Data FSSig. 24a)^50^. Furthermore, given the lack of mechanistic insight provided by conventional analysis methods, any observed difference could reasonably have been interpreted as evidence that apoptosis makes a partial and/or negligible contribution to the lethality of transcriptional inhibition. Our initial evaluations of transcriptional inhibitors used the drug GRADE analysis method^51^, and the mechanistic clarity provided by GRADE-based analysis revealed that apoptotic-deficient cells completely arrest cell proliferation without activating any cell death (Extended Data FSSig. 24b).

Our study also identifies two key regulators of this process: PTBP1 and BCL2L12. PTBP1 is a ubiquitously expressed RNA binding protein with several established roles in RNA and translational regulation^38,40^. Notably, these prior functions appear not to contribute to lethality following Pol II degradation. Our study, however, confirms that PTBP1 can translocate to the cytoplasm in response to stress, which in the context of Pol II degradation, appears to communicate information about Pol II protein level from the nucleus to the cytoplasm, facilitating the activation of apoptosis. Additionally, *BCL2L12* is also a ubiquitously and highly expressed gene. This feature distinguishes *BCL2L12* from other members of the BCL2 family, which generally have low and varied expression across cell types. Furthermore, BCL2L12 is generally considered an anti-apoptotic protein, and has not been described to have any pro-apoptotic functions^52^. Thus, it appears that PTBP1 and BCL2L12 have unique roles in response to Pol II degradation that are distinct from their previously annotated functions. Central remaining questions include developing a detailed molecular mechanism by which these proteins activate apoptosis following Pol II degradation, and exploring the context in which this pathway evolved to be beneficial.

Prior to this discovery, substantial attention had already been focused on targeting the transcriptional cycle in the context of cancer treatment. New drugs that target transcriptional Cdk proteins appear to be promising. However, the mechanisms allowing for the selective killing of cancer cells remain unclear, and are generally presumed to be dependent on loss of rapidly turned over proteins, such as Myc or Mcl-1^53^. Our study highlights that many of these drugs owe their lethality to the Pol II degradation-dependent mechanism that we describe. Notably, the definitions used in this study for defining a Pol II degradation-dependent cell death are conservative, as a drug was only characterized as PDAR-dependent if no other backup mechanisms existed to facilitate death in the absence of this pathway. Importantly, our study also highlights that selective killing is possible for PDAR-dependent drugs, as we identified several conventional DNA damaging chemotherapies, which have long-standing clinical utility, that also unexpectedly score as PDAR-dependent using our conservative criteria. Thus, future studies of the PDAR pathway are likely to reveal new biomarkers that improve our ability to use a wide array of anti-cancer drugs, and also may reveal new ways to selectively kill cancer cells.

## Acknowledgements

We thank M. Walhout, T. Fazzio, E. Baehrecke, R. Davis, C. Navarro, T. Naylor, M. Wesley, and all members of the Lee Lab and UMass Chan Medical School DSB community for their helpful comments and suggestions throughout the development and execution of this study. We especially thank E. Calvo-Roitberg for advice related to evaluating PTBP1-dependent splicing and RNA location. Additionally, we thank J. Dekker for providing the HAP1-RPB1-AID cells.

## Author contributions

N.W.H. and M.J.L. conceptualized this project, including selection of methodology, analytical strategies, data visualization, and interpretation. N.W.H. performed all experiments and analyses in this study. G.A.B. assisted N.W.H. with the evaluation of cell death mechanisms activated by transcriptional inhibitors and canonical cell death activating drugs. M.E.H. generated the materials used in the chemo-genetic profiling of triptolide, and assisted with the design and analysis of chemo-genetic profiling data. A.A.P. designed the experiments and analytical strategy used in evaluating PTBP1- and triptolide-dependent changes in RNA localization and RNA splicing. The original draft of this manuscript was written by N.W.H. and M.J.L. All authors participated in manuscript review and editing.

## Data and materials availability

RNA sequencing data generated in this study can be found as FASTQ files from the Gene Expression Omnibus (GEO) with the following accession numbers: GSE283148 (triptolide exposure time course in U2OS and BAX/BAK DKO cells), GSE283149 (triptolide exposure for 8 hours in NUP93 or EXOSC5 KO cells), GSE283150 (RNA sequencing from nuclear and cytoplasmic fractionation in U2OS or PTBP1 KO cells), GSE283151 (triptolide exposure following a-amanitin in Pol II switchover system). Chemo-genetic profiling data can be found as raw FASTQ files (GSE283147), and as sequencing counts or death rates in Supplementary Tables 3 and 4, respectively. Source data associated with Fig. 1e can be found in Supplementary Table 1. Source data associated with Fig. 2 can be found in Supplementary Table 2. Source data associated with Fig. 5c can be found in Supplementary Table 5. Source data associated with Fig. 5g can be found in Supplementary Table 6. All other data are available in the main text or supplementary materials.

## Code Availability

Custom MATLAB scripts for dose-response curve fitting, lethal fraction kinetics, and drug GRADE computation are deposited on GitHub (https://github.com/MJLee-Lab). Custom scripts for all other analyses are available upon request.

## Materials & Correspondence

Correspondence and requests for materials should be directed to Michael Lee.

## MATERIALS AND METHODS

### Cell lines and reagents

U2OS, HT-29, H1650, WI-38, Hs578T, MDA-MB-453, MCF7 and MDA-MB-157 cells were obtained from the American Type Culture Collection (ATCC). A375 cells were a gift from the Green laboratory (UMass Chan Medical School). HAP1 and HAP1-RPB1-AID cells were a gift from the Dekker laboratory (UMass Chan Medical School)^19^. RPE-1 cells were a gift from the Pazour laboratory (UMass Chan Medical School). PC9 cells were a gift from the Pritchard laboratory (Penn State University). U2OS, MDA-MB-157, A375, Hs578T, MDA-MB-453, and MCF7 cells were grown in DMEM (Corning, 10-017-CV) supplemented with 2mM glutamine (Corning, 25-005-CI). HT-29 cells were grown in McCoy’s 5A medium (Corning, 10-050-CV). WI-38 cells were grown in MEM medium (Thermo Fisher Scientific, 10370088). H1650 and PC9 cells were grown in RPMI medium (Thermo Fisher Scientific, 11875119). RPE-1 cells were grown in low glucose DMEM (Gibco, 11885-092) and Hams F-12 (Thermo Fisher Scientific, 11765054) mixed at a 1:1 ratio. HAP1 cells were grown in IMDM medium (Thermo Fisher Scientific, 12440053). Each media was supplemented with 10% FBS (Peak Serum, PS-FB2, lot no. 21E1202) and penicillin-streptomycin (Corning, 30-002-CI). Hs578T cells were supplemented with 10µg/mL insulin (Thermo Fisher Scientific, 12585014). Cell lines were cultured in incubators at 37C with 5% CO2. Cell lines were maintained at low (<25) passage number from the original vial.

U2OS-BAX/BAK1 double knockout cells (DKO) were generated previously^54^. All mKate2+ cells were generated by lentiviral infection with NucLight Red Lentivirus (Sartorius, 4627), followed by FACS selection. HAP1-RPB1-AID cells were treated with 450 µg/mL hygromycin for one week prior to use. U2OS-Cas9 expressing and U2OS-TP53 knockout cells were generated previously^34^.

U2OS-PTBP1 and U2OS-BCL2L12 clonal knockout cells were generated using CRISPR. For each gene, the highest-scoring sgRNA was selected from the TKOv3 library (PTBP1: 5’ – CGAGGTGTAGTAGTTCACCA – 3’; BCL2L12: 5’ – AGAAGGAAGCCATACTGCGG – 3’) and cloned into the pX330-puro plasmid using the single-step digestion-ligation protocol from the Zhang laboratory (available on the Zhang laboratory Addgene page). U2OS cells were then transiently transfected with the respective pX330-sgRNA-puro constructs using the FuGENE HD Transfection Reagent (Promega, E2311). Transfected cells were then selected with 1 µg/mL puromycin for 3 days, followed by replating and recovery for 2 days. Single cell clones were generated using limiting dilution. Clones were validated by sequencing, phenotyping, and for PTBP1 knockouts, immunoblotting.

U2OS-MCL1, U2OS-NUP93 and U2OS-EXOSC5 knockout cells were generated using a transient transfection strategy. Briefly, sgRNA (MCL1: 5’ – CATGTAGAGGACCTAGAAGG – 3’; NUP93: 5’ – GAACTTACAGGAGATCCAGC – 3’; EXOSC5: 5’ – GAAGTTCTTACCTTGCAGGA – 3’) were cloned into pX330-puro as described above. U2OS cells were plated in six-well plates (300,000 cells per well) and left to adhere overnight. The following day, cells were transiently transfected with 2 µg of respective pX330-sgRNA-puro construct using FuGENE HD Transfection Reagent. Cells were then selected using 1 µg/mL puromycin for either 2 days (for NUP93 and EXOSC5) or 3 days (for MCL1), followed by replating and expansion for either 1 day (for NUP93 and EXOSC5) or 2 days (for MCL1) prior to seeding for subsequent assays. Control cells were similarly transfected with non-targeting sgRNA (NONT: 5’ – CGGCGGTCACGAACTCCAGC – 3’).

For the RPB1 switchover system, a FLAG-RPB1-N792D-ΔCTD fragment was PCR amplified (Forward primer: 5’ – CAATTCCACAACACTTTTGTCTTATACTTGGATCCATGGACTACAAGGACGACGATGACA - 3’; Reverse primer: 5’ – TAGGGGGGGGGGAGGGAGAGGGGCCGGCCGGGGCTCAGCTGGGAGACATGGCACCAC – 3’) from FLAG-Pol2-WT (Addgene, 35175)^55^. The PCR fragment resulted in a RPB1 mutant that lacked the entire C-terminal domain (repeats 1-52) and included an N-terminal FLAG tag, an ⍺-amanitin resistance mutation, and homology regions to facilitate insertion into pLVX-Tre3G-IRES (Clontech, 631362)—a lentiviral vector for doxycycline-inducible gene expression. pLVX-Tre3G-IRES was digested with BamHI and NotI, gel extracted and purified. The PCR product was then inserted into the MCS1 of pLVX-Tre3G-IRES using Gibson Assembly (New England Biolabs, E2611). The resulting plasmid, pLVX-Tre3G-FLAG-RPB1-N792D-ΔCTD, was packaged into virus using Lenti-X 293T cells (Takara, 632180), along with a vector expressing the Tet-On 3G transactivator protein, pLVX-Tet3G (Clontech, 631358). U2OS cells were infected with pLVX-Tet3G lentivirus, followed by selection with 600 µg/mL G418. Tet-3G expressing cells were then infected with pLVX-Tre3G-FLAG-RPB1-N792D-ΔCTD virus, and selected with 1 µg/mL puromycin. Single cell clones were generated using limiting dilution. Clones were selected for transgene expression levels approximating endogenous Rpb1 levels, determined using immunoblotting.

Compounds used in this study were as follows. SYTOX Green (Thermo Fisher Scientific, S7020), Triptolide (Sigma-Aldrich, T3652), ⍺-amanitin (Sigma-Aldrich, A2263), ABT-737 (ApexBio, A8193), RSL3 (ApexBio, B6095), Elesclomol (Sigma-Aldrich, SML2651), MNNG (Biosynth Carbosynth, FM11256), TNF-⍺ (ApexBio, P1001), SM-164 (ApexBio, A8815), IAA (Sigma-Aldrich, I5148), z-VAD-FMK (ApexBio, A1902), VX765 (ApexBio, A8238), TTM (Sigma-Aldrich, 323446), Rucaparib (ApexBio, A4156), Necrostatin-1 (SelleckChemicals, S8037), Ferrostatin-1 (SelleckChemicals, S7243), Doxycycline Hyclate (Sigma-Aldrich, D5207), Actinomycin-D (SelleckChemicals, S8964, Ethynylcytidine (MedChemExpress, HY-16200), Cordycepin (Sigma-Aldrich, C3394), Flavopiridol (SelleckChemicals, S1230), DRB (Sigma-Aldrich, D1916), THZ1 (SelleckChemicals, S7549), THZ531 (MedChemExpress, HY-103618), YKL-5-124 (MedChemExpress, HY-101257B), Abemaciclib (MedChemExpress, HY-16297A), Palbociclib (ApexBio, A8316), Ribociclib (MedChemExpress, HY-15777), PF3600 (SelleckChemicals, S8816), PF4091 (ChemieTek, CT-PF0710), 4-NQO (SelleckChemicals, E0155), Cisplatin (SelleckChemicals, S1166), Carboplatin (Sigma-Aldrich, C2538), Camptothecin (SelleckChemicals, S1288), Topotecan HCl (SelleckChemicals, B2296), Idarubicin HCl (SelleckChemicals, S1228), Etoposide (SelleckChemicals, S1225), Teniposide (S1787 SelleckChemicals), Niraparib (SelleckChemicals, S2741), Rucaparib phosphate (SelleckChemicals, S1098), Cycloheximide (Sigma-Aldrich, C1988), Homoharringtonine (Sigma-Aldrich, SML1091), Anisomycin (Sigma-Aldrich, A9789), Bortezomib (ApexBio, A2614), MG-132 (SelleckChemicals, S2619), Brefeldin A (Sigma-Aldrich, B7651), Thapsigargin (Sigma-Aldrich, T9033), Tunicamycin (Sigma-Aldrich, T7765), ABT-199 (ApexBio, A8194), ABT-263 (ApexBio, A3007), Panobinostat (ApexBio, A8178), Vorinostat (ApexBio, A4084), Torin 2 (ApexBio, B1640), Rapamycin (ApexBio, A8167), Mirdametinib (MedChemExpress, HY-10254), Buparlisib (MedChemExpress, HY-70063), Capivasertib (MedChemExpress, HY-15431), SGI-1027 (ApexBio, B1622), JQ1 (SelleckChemicals, S7110), Staurosporine (SelleckChemicals, S1421), H2O2 (Fisher Scientific, BP2633500), LPS (Sigma-Aldrich, L4391), Nigericin (ApexBio, B7644), copper(II) chloride (ThermoFisher Scientific, 405845000), S63845 (SelleckChemicals, S8383).

Primary antibodies used in this study were as follows. Rpb1-NTD rabbit mAb (Cell Signaling Technology, 14958, 1:1,000 dilution), β-actin mouse mAb (Sigma-Aldrich, A2228, 1:15,000 dilution), DYKDDDDK Tag (FLAG) rabbit mAb (Cell Signaling Technology, 14793S, 1:1,000 dilution), PTBP1 rabbit mAb (Cell Signaling Technology, 57246S, 1:1,000 dilution), Histone-H3 rabbit mAb (Cell Signaling Technology, 4499S, 1:2,000 dilution), ⍺-tubulin mouse mAb (Sigma-Aldrich, T6199, 1:10,000 dilution), pH2A.X (Ser139) rabbit mAb (Cell Signaling Technology, 9718, 1:1,000 dilution), Mcl-1 rabbit mAb (Cell Signaling Technology, 5453, 1:1,000 dilution), Apaf-1 rabbit mAb (Cell Signaling Technology, 8723S, 1:1,000 dilution).

### Immunoblotting

Cells were seeded in 6-well plates (300,000 cells per well) or 10-cm dishes (1.5 x 10^6^ cells per dish) and left to adhere overnight, unless otherwise stated. Cells were treated the next morning. At indicated timepoints, media were removed and collected. Samples were washed once with PBS, and the wash was added to the collected media. SDS-lysis buffer (50 mM Tris-HCl, 2% SDS, 5% glycerol, 5 mM EDTA, 1 mM NaF, 10 mM β-glycerophosphate, 1 mM PMSF, 1 mM Na_3_VO_4_, protease inhibitor tablet and phosphatase inhibitor tablet) was used to lyse the cells, along with any dead cells that were pelleted from the media and wash. Lysates were centrifuged through an AcroPrep 96 well 3.0 µm glass fiber/0.2 µm Bio-Inert membrane filter plate (Pall Laboratory, 5053). Protein content was quantified with a BCA assay (Thermo Fisher Scientific, 23225) and normalized for equal loading. Samples were boiled for 5 minutes at 95C in 6x Laemmli buffer. Denatured samples were run on hand-poured SDS-PAGE gels. Gels were subsequently wet transferred onto nitrocellulose membranes and blocked in 1:1 PBS: Intercept Blocking Buffer (LI-COR, 927-70003) for 1 hour at room temperature. Membranes were then incubated overnight on a rocking shaker at 4C in primary antibody diluted in 1:1 PBS-0.1% Tween: Intercept Blocking Buffer. The following day, membranes were incubated in primary anti–β-actin antibody (diluted 1:15,000) for 1 hour at room temperature, followed by two 5-minute washes with PBS-0.1% Tween. Membranes were then incubated with secondary antibodies (diluted 1:15,000 in 1:1 PBS-0.1% Tween: Intercept Blocking Buffer) conjugated to infrared dyes (LI-COR, IRDye 680RD, goat anti-mouse immunoglobulin G (IgG) secondary, catalog no. 926-68070; IRDye 800CW, goat anti-rabbit IgG secondary, catalog no. 926-32211) for 1 hour at room temperature. Following four 5-minute washes with PBS-0.1% Tween and one 5-minute wash with PBS, blots were visualized using a LI-COR Odyssey CLx scanner. Raw protein expression levels were quantified in LI-COR’s ImageStudio software, background subtracted, divided by β-actin signals to normalize for loading differences, then normalized per gel to a reference sample (generally, an untreated sample).

### Proteome profiler-based analysis of apoptotic proteins

Proteome Profiler Apoptosis Arrays (R&D Systems, ARY009) were performed according to the manufacturer’s instructions. Briefly, U2OS cells were collected at 1.5 million cells per condition and lysed according to instructions. An equal volume of lysate (250 µL) was loaded per array. Blot arrays were visualized using a LI-COR Odyssey CLx scanner, and protein expression was quantified in LI-COR’s ImageStudio software.

### Labelling and measurement of nascent transcription

Methods for labelling and quantification of nascent transcription were adapted from methods described previously^56^. Cells treated with transcriptional inhibitors were labeled with 1 mM 5-ethynyl uridine (Sigma-Aldrich, 909475) for 1 hour in existing media. Cells were then washed with PBS, trypsinized, pelleted, and washed with cold PBS. Cells were then fixed in 4% fresh formaldehyde in PBS at room temperature for 15 minutes. Fixed cells were washed with cold PBS, pelleted, resuspended in ice-cold 100% methanol, and stored at -20C overnight. The following day, the methanol was removed, and the cells were washed twice with PBS-0.1% Tween. Cells were then incubated for 30 minutes at room temperature with 86.5 µL Click-iT reaction buffer, 4 µL CuSO_4_ buffer, 0.125 µL Alexa Fluor-azide-488 and 10.3 µL Click-iT reaction buffer additive (all from Thermo Fisher Scientific, C10329). Cells were washed once with 100 µL Click-iT reaction rinse buffer (from Thermo Fisher Scientific, C10329), then washed two more times with PBS-0.1% Tween. Samples were resuspended in PBS-0.1% Tween, filtered, and run on a Miltenyi MACSQuant VYB cytometer with laser and filter settings appropriate for reading Alexa-488. Data were analyzed using FlowJo software.

### RNA sequencing with ERCC spike-ins for absolute RNA quantification

For evaluation of mRNA abundance following long-term triptolide exposure in DKO cells, DKO and WT cells were seeded onto six-well plates (300,000 cells per well) and left to adhere overnight. Cells were then drugged the next day (D0). To evaluate the transcriptome of WT cells after 1 day of triptolide, cells were cotreated with 50 µM z-VAD to inhibit loss of material from cell death. Due to the long assay length, plating and drugging was staggered for each condition, such that the assay endpoint for each sample occurred simultaneously. For analyses that defined the 50 shortest or 50 longest half-life RNAs, these lists of RNAs were based on prior studies^57^.

For evaluation of mRNA abundance in the context of NUP93 or EXOSC5 knockout, knockout and nontargeting cells were seeded onto six-well plates (300,000 cells per well) and left to adhere overnight. Cells were then drugged the next day.

For evaluation of mRNA abundance in Pol II switchover cells, Rpb1-N792D-ΔCTD expressing cells were seeded onto six-well plates (100,000 cells per well) and left to adhere overnight. The following day, the media was removed and replaced with fresh media with or without 2 µg/mL doxycycline. 48 hours later, drug or vehicle was gently added to the existing media.

For the extraction of mRNA for the experiments above, at assay endpoint cells were trypsinized, counted using a hemacytometer, pelleted, washed with PBS, and moved immediately to RNA extraction. Total RNA was extracted using the RNeasy Plus Mini Kit (Qiagen, 74134) according to manufacturer protocol. To facilitate absolute RNA quantification, polyadenylated ERCC standards (Thermo Fisher Scientific, 4456740) were diluted 1:100 in nuclease-free water, and added to each sample at 1 µL/100,000 cells. ERCC spike-ins were added immediately following lysis of cells in RLT-plus buffer, and prior to all subsequent total RNA extraction steps using the RNeasy Plus kit. All RNA integrity numbers (RIN) were greater than 9, as measured on a 4150 Tapestation System (Agilent Technologies). mRNA was isolated using the NEBNext Poly(A) mRNA Magnetic Isolation Module (New England Biolabs, E7490). RNAseq libraries were prepared with the NEBNext Ultra II Directional RNA Library Prep Kit using the NEBNext Multiplex Oligos for Illumina (New England Biolabs, E7765 and E7600). Paired-end 2 x 50nt sequencing of libraries was performed in-house on the Illumina NextSeq2000 using the NextSeq 1000/2000 P2 XLEAP-SBS Reagent Kit (Illumina, 20100987).

Quality control for the RNAseq datasets were performed using FastQC. Transcript abundance was estimated using Kallisto (0.46.1) with parameters --bootstrap-samples 30 -- single-overhang --rf-stranded^58^. The Ensembl GRCh38 cDNA transcriptome build, appended with ERCC control transcript sequences, was used to build a Kallisto index. Gene-level counts were generated in R using the tximport package^59^. Counts were normalized to ERCC spike-ins using DESeq2, implemented with the function “estimateSizeFactors” with the option “controlGenes” set to the identities of the 92 ERCC transcripts^60^. Counts were filtered for protein coding genes in R using AnnotationHub. Differential gene expression was performed with DESeq2, and fold change shrinkage was performed using the function lfcShrink with the adaptive shrinkage estimator “ashr”^61^.

### Extraction of RNA and protein from nuclear and cytoplasmic fractions

Nuclear-cytoplasmic fractionation was performed in such a way to facilitate extraction of RNA and protein from the same cellular lysate. A similar protocol, excluding the RNA extraction steps, was used in fractionation experiments that only examined protein expression. Cells were plated at 6 million cells per condition (3 million cells per 15 cm plate) and left to adhere overnight. Cells were then drugged the next day. At assay endpoint cells were trypsinized, counted using a hemacytometer, pelleted, washed with PBS, and moved immediately to cellular fractionation. Cells were first resuspended in 900 µL hypotonic lysis buffer (50 mM HEPES pH 7.2, 150mM NaCl, 0.5 mM EDTA, 0.15% Triton X-100, 1 mM PMSF, protease inhibitor tablet, phosphatase inhibitor tablet, 20 U/mL SUPERase-In RNase inhibitor (Thermo Fisher Scientific, AM2694)). Cells were pipet up and down 10 times, 1/3 of the lysate was taken as a whole-cell lysate (WCL) sample, and the cells were left to swell on ice for 10 minutes. Samples were centrifuged at 12,000xg for 30 seconds at 4C. The supernatant was collected as the cytoplasmic fraction (CYT). The nuclear pellet was washed once with 500 µL wash buffer (PBS containing 0.1% Triton X-100, 1mM EDTA and 20 U/mL SUPERase-In), centrifuged at 12,000xg for 30 seconds at 4C, and the wash buffer was completely removed. The nuclear pellet was then lysed in 400µL 1x nuclear lysis buffer (50 mM HEPES pH 7.2, 150mM NaCl, 0.5 mM EDTA, 1% Triton X-100, 0.5% Sodium Deoxycholate, 0.2% SDS, 1 mM PMSF, protease inhibitor tablet, phosphatase inhibitor tablet, 20 U/mL SUPERase-In). 10x nuclear lysis buffer was added to WCL and CYT fractions to equalize surfactant amounts to that of the nuclear fraction. Each resulting lysate was divided in half to facilitate either downstream protein analysis or RNA extraction.

For protein fractions, 100x Benzonase buffer (50 mM Tris-HCl, 20 mM NaCl, 2 mM MgCl_2_, 2000 U/mL Benzonase Nuclease (Sigma Aldrich, E1014)) was added to lysates. Lysates were kept on ice for 30 minutes with periodic vortexing. The BCA assay was performed on all cellular fractions, and 10 µg of each lysate was subjected to immunoblotting as described above. Total protein was detected using the Revert 700 Total Protein Stain Kit (LI-COR, 926-11010) prior to incubation with primary antibody.

For extraction of total RNA from RNA fractions, samples were subjected to the RNeasy Plus manufacturer protocol with the following modifications. 3.5x sample volume of RLT-plus buffer was added to cellular fractions. Prior to removal of genomic DNA using the gDNA Eliminator spin column, ERCC standards (diluted 1:10) were added to each lysate at a concentration of 1 µL/ 1 million cells (or similarly, 1 million nuclei or 1 million cell cytoplasm’s). Following genomic DNA removal, 2.5x original lysate volume of 100% ethanol was added to each flowthrough, and subsequent steps of the RNeasy Plus protocol were followed as standard. mRNA was isolated as above. Sequencing libraries were prepared with the NEBNext Ultra II Directional RNA Library Prep Kit, using manufacturer recommended modifications to generate large (∼450nt) inserts. Paired-end 2 x 150nt sequencing of libraries was performed in-house on the Illumina NextSeq2000 using the NextSeq 2000 P3 Reagent Kit (Illumina, 20040561).

### Alternative splicing analysis from RNA sequencing data

To facilitate detection of splice-level events, cytoplasmic and nuclear RNAseq FASTQ files corresponding to the same biological samples (above) were pooled. Regulated splicing events were detected and analyzed with VAST-TOOLS^62,63^. Briefly, reads were aligned to the human genome (hg38) using the align function. Differential splicing events were identified using the diff function (with parameters -S 2). Events were filtered for adequate coverage using the tidy function (with parameters -min_N 4, --noVLOW, --min_SD 5, and --p_IR). Splicing events were considered significantly different if the VAST-TOOLS reported expected absolute ΔPSI value was greater than 0.1 and if the minimum value (95% confidence) of the absolute ΔPSI value was greater than 0.

### STACK assay for kinetic evaluation of drug responses

The STACK assay was used to evaluate cell proliferation and cell death continuously over time using microscopy, as previously described^13^. Briefly, mKate2+ cells were seeded into 96-well black-sided plates, and left to adhere overnight. The following day, cells were given drugged media containing 50 nM SYTOX Green, and imaged over time time-lapsed images were collected using an IncuCyte S3 (Essen Biosciences). Images were collected with a 10x objective with acquisition in the green channel ex: 460 ± 20, em: 524 ± 20, acquisition time: 300ms; and red channel ex:585 ± 20, em: 635 ± 70, acquisition time: 400ms. Counts of dead (green) and live (red) cells were determined using the built-in IncuCyte software (Essen Biosciences) and exported for analysis using a custom MATLAB script.

For the long-term evaluation of BAX/BAK DKO cells following triptolide, accurate dead cell measurements using STACK as described above are insufficient, as identification of a dead cell from an image is no longer possible after the dead cell fully degrades away. To overcome this technical confound, a modified STACK assay was used, taking advantage of the FLICK assay (described in detail below), which does not rely on image segmentation for cell quantification. Cells were seeded and drugged as above, at a high density (15,000 cells per well). Separate plates were prepared for each timepoint, whereby cells were first imaged using an EVOS FL Auto 2 automated microscope (ThermoFisher Scientific). Images were acquired using a 10x objective (EVOS 10x objective, Cat #: AMEP4681). Sytox images were acquired using a GFP filter cube (EVOS LED Cube, GFP, Cat #: AMEP4651, ex: 470/22, em: 525/50, acquisition time: 13.5ms) Mkate2+ images were acquired using a TexasRed filter cube (EVOS LED Cube TxRed, Cat #: AMEP4655, ex: 585/29, em: 628/32, acquisition time: 642.0ms). Following imaging, FLICK was performed as an endpoint measurement to obtain lethal fraction values, as described below. Images were linearly adjusted using a custom MATLAB script.

### Quantification of drug-induced cell death using the FLICK assay

The FLICK assay was performed as described^54,64^. Briefly, 90 µL of cells were seeded at a density of 2,000-5,000 cells per well in black-sided optical-bottom 96-well plates (Greiner Bio-One, 655090) and left to adhere overnight. The following day, cells were treated with 10 µL of media containing the indicated compound/s along with SYTOX Green, resulting in a final SYTOX concentration of 2 µM. Fluorescence was monitored at various times following drugging with a Tecan Spark (running SparkControl software version 2.2) microplate reader (ex: 503, em: 524). Gain was set to achieve a linear relationship between SYTOX signal and dead cell number. To obtain the total cell number at the time of drugging, a duplicate “T0” plate was lysed in 0.15% Triton X-100 and 2 µM SYTOX for 2-4 hours at 37C prior to taking a fluorescence reading. Total cell number for each condition at the end of the assay was similarly determined by lysing cells in 0.15% Triton X-100. From these measurements (dead cells at each timepoint, along with total cell numbers at the beginning and end of the assay), live cell numbers over time can be inferred and all calculations to determine LF, FV, RV and GR can be made ^50^:

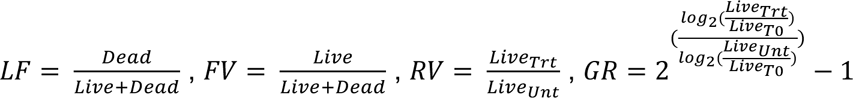

All analyses were performed using custom MATLAB scripts. Dose-response curves were fit using a 4-parameter sigmoidal equation. Lethal fraction kinetics were fit using the lag-exponential death (LED) equation ^13^:

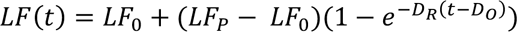

All analyses were performed using custom MATLAB scripts. All data were fit with the “fit” function in MATLAB using nonlinear least-squares.

### Drug GRADE-based analysis of drug responses

True proliferation rates and death rates were calculated using the GRADE method, as previously described^51^. Conceptually, the GRADE method infers the true, underlying cell proliferation rates and death rates of cell population by cross-referencing experimentally measured death rates (FV or LF) and net population growth rates (GR) with computational simulations. To begin, we simulated all pairwise combinations of 500 proliferation rates and 500 death rates using the following equations where *C_0_* = initial cell number, *t* = assay length, *τ* = proliferation rate, and *D_R_* = death rate.

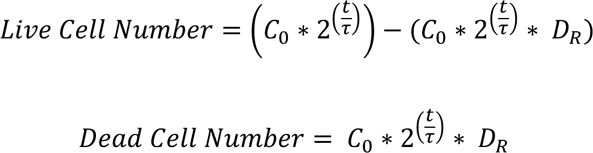

FV and GR values (defined above) were then calculated for each simulated proliferation rate and death rate pair. Next, for a given condition, experimentally determined FV and GR values were calculated (as above). The simulated proliferation rate and death rate that yield the empirically determined FV and GR values was then found.

### Quantification of Pol II half-life

Half-lives for Pol II-a and Pol II-o were determined using temporally resolved quantitating immunoblotting using an antibody recognizing the N terminal domain of Rpb1 (Cell Signaling, 14958S), as described above. Immunoblots were quantified in ImageStudio 3.1.4, normalized to β-actin, then normalized to the untreated sample (time 0) on each gel respectively, generating an abundance value relative to steady state. Data were fit to a two-term exponential decay function in MATLAB, relative to the highest Rpb1 signal observed over the time course. Half-lives determined from the fits, as the time at which half-maximal signal was reached. To correlate Pol II half-lives with cell death timing, the LF50 time was similarly found from lag-exponential death (LED) fits to STACK data. LF50 was defined as the time at which half of the maximum observed LF (measured or inferred after 72 hours) was reached.

### Pol II Switch over system

For experiments involving the RPB1 switchover system, cells were seeded as described for the respective experiment. The following day, cells we treated with or without 2 µg/mL Doxycycline (Sigma, D5207) for 48 hours to induce the expression of FLAG-RPB1-N792D-ΔCTD. 10 µM ⍺-amanitin was added to the cells without removing the conditioned media to degrade the endogenous Pol II.

### Chemo-genetic profiling of triptolide-induced cell death

Conventional approaches used for these “chemo-genetic profiling” studies are unable to accurately identify death-regulatory genes due to the confounding effects of varied proliferation rates between cells harboring different gene knockouts^34^. In general, specialized analytical methods are required to identify death-regulatory genes when drug-induced lethality occurs in a proliferative population, at low rates relative to proliferation, or via non-apoptotic mechanisms. However, in this case, we recognized that triptolide-induced death is: 1) exclusively apoptotic, 2) occurring rapidly (before the effects of varied proliferation can be observed in the population), and 3) activated at high rates (such that large numbers of dead cells accumulate before apoptotic corpses have decayed). Thus, we developed a simple experimental approach to mechanically separate live cells from apoptotic corpses, such that both populations could be sequenced independently.

The genome-wide CRISPR screen was performed using the Toronto KnockOut Library v3 (TKOv3), which contains 4 sgRNA per gene, along with 142 non-targeting sgRNA^65^. U2OS-Cas9 expressing cells were selected with 5 µg/mL blasticidin for 5 days. Following selection, cells were divided into two replicates, which would be handled separately for the remainder of the assay. Cells were then transduced with the TKOv3 viral library using a ‘spinfection’ method. To maintain sufficient library coverage, more than 200 million cells were transduced per replicate, at an MOI of 0.3. To do this, cells were divided into 12-well plates at a density of 2 million cells per along with 75 µL virus and 0.8 µg/mL polybrene (Millipore, TR1003G). Plates were centrifuged at 37 °C for 2 h at 830*g*. Following spinfection, virus-containing media was gently removed and replaced with fresh media, and cells were returned to the incubator overnight. The following morning, cells were moved out of the 12-well plates and onto 15 cm dishes. The following day, cells were treated with 1 µg/mL puromycin for 3 days, followed by a two-day expansion without selection. Cells were then seeded onto 15cm plates and let to adhere overnight prior to drugging. At the time of drugging, a T0 sample was taken from the cell population and frozen, at a population size 650x that of the library size. The rest of the cells were treated with either 1 µM triptolide or DMSO. Cells were incubated for 32 hours in drug, resulting in approximately 50% cell death. Apoptotic corpses, which become non-adherent, were removed and saved. Plates were washed with PBS to dislodge any remaining dead cells, and PBS wash was added with the separated dead cell fraction. The adherent cells remaining on the plate were the live cell population. The live cells were trypsinized and collected. Each condition was collected such that a minimum of 650x library coverage was achieved per replicate.

Genomic DNA was extracted using the Wizard Genomic DNA Purification Kit (Fisher Scientific, PR-A1125). sgRNA sequences were PCR amplified out of the genomic DNA, and gel-extracted and purified. A second round of PCR added sequencing adaptors and multiplexing barcodes, and libraries were pooled and sequenced on a HiSeq 4000 at 500× coverage.

The FASTX-Toolkit (0.0.14) was used to trim reads to get just CRISPR guide sequences (parameters: -f 23 -l 42). Trimmed reads were mapped to the TKOv3 library with Bowtie (1.3.0) allowing for 2 mismatches (parameters: -v 2 -m 1). Samples were normalized for sequencing depth using median of ratios method implemented in DESeq2. Guides with low counts (approximately the bottom 2% of guides) were removed from the library. sgRNA-level log_2_ fold-changes were determined using DESeq2, and were then z-scored to the distribution of non-targeting sgRNA. The median sgRNA-level z-score was determined for each gene to collapse scores to the gene level. An empiric *P* value was determined for each gene by bootstrapping from the sgRNA-level fold change scores of non-targeting guides, and was FDR corrected using the Benjamini–Hochberg procedure. An FDR cutoff of 0.1 was used to determine genes that significantly alter the cell death rate following triptolide. Gene set enrichment using an FDR-corrected one-tailed Fishers Exact test was performed to test if any apoptotic-related pathways from the Molecular Signatures Database (MSigDB) were overrepresented among genes whose knockout significantly reduced triptolide-induced death.

### Evaluation of drug TIS score

FLICK was used to determine lethal fraction kinetics in U2OS, PTBP1-KO, BCL2L12-KO, and *BAX/BAK* DKO cells across a panel of 46 drugs, across 7 half-log dilutions, and spanning the following 16 classes. Pol II degraders (Triptolide, 1 – 0.001 µM; ⍺-amanitin, 10 – 0.01 µM). RNA synthesis inhibitors (Actinomycin-D, 1 – 0.001 µM; Ethynylcytidine, 10 – 0.01 µM; Cordycepin, 100 – 0.1 µM). Transcriptional CDK inhibitors (Flavopiridol, 10 – 0.01 µM; DRB, 100 – 0.1 µM; THZ1, 10 – 0.01 µM; THZ531, 10 – 0.01 µM; YKL-5-124, 31.6 – 0.0316 µM). Cell cycle CDK inhibitors (Abemaciclib, 10 – 0.01 µM; Palbociclib, 10 – 0.01 µM; Ribociclib, 10 – 0.01 µM; PF3600, 31.6 – 0.0316 µM; PF4091, 31.6 – 0.0316 µM). DNA crosslinking agents (4-NQO, 31.6 – 0.0316 µM; Cisplatin, 100 – 0.1 µM; Carboplatin, 316 – 0.316 µM). Topoisomerase I inhibitors (Camptothecin, 10 – 0.01 µM; Topotecan, 31.6 – 0.0316 µM). Topoisomerase II inhibitors (Idarubicin, 1 – 0.001 µM; Etoposide, 31.6 – 0.0316 µM; Teniposide, 31.6 – 0.0316 µM). PARP inhibitors (Niraparib, 100 – 0.1 µM; Rucaparib, 100 – 0.1 µM). Translational inhibitors (Cycloheximide, 100 – 0.1 µM; Homoharringtonine, 10 – 0.01 µM; Anisomycin 316 – 0.316 µM). Proteasome inhibitors (Bortezomib, 1 – 0.001 µM; MG132, 10 – 0.01 µM). ER- and proteotoxic-stress inducers (Brefeldin A, 31.6 – 0.0316 µM; Thapsigargin, 31.6 – 0.0316 µM; Tunicamycin, 10 – 0.01 µM). BH3 mimetics (ABT-199, 100 – 0.1 µM; ABT-263, 100 – 0.1 µM; ABT-737, 31.6 – 0.0316 µM). HDAC inhibitors (Panobinostat, 1 – 0.001 µM; Vorinostat, 100 – 0.1 µM). mTOR inhibitors (Torin2, 31.6 – 0.0316 µM; Rapamycin, 10 – 0.01 µM). MEK/PI3K/AKT inhibitors (Mirdametinib, 31.6 – 0.0316 µM; Buparlisib, 31 – 0.0316 µM; Capivasertib, 31.6 – 0.0316 µM). Others (SGI-1027, 31.6 – 0.0316 µM; JQ1, 100 – 0.1 µM; Staurosporine, 10 – 0.01 µM).

Lethal fraction kinetics were fit to a lag exponential death (LED) equation. Non-lethal conditions were defined as those having a maximum observed lethal fraction less than 0.16, approximately double the death observed in untreated cells. Cells were further classified as apoptotic or non-apoptotic by thresholding a 50% reduction in maximum lethal fraction in the BAX/BAK DKO background compared to U2OS cells. To determine the “Transcription Inhibition Similarity” (TIS) score for a particular drug-dose pair, the area under the LED fit (AUC) was determined, and the baseline cell death was removed by subtracting out the AUC of untreated cells. The difference in drug response between KO and U2OS cells was defined as 1 minus the AUC ratio of KO over U2OS. This value, determined for PTBP1-KO and BCL2L12-KO separately, was summed. Finally, scores were divided by that of 1 µM triptolide, such that a score of one denotes a functional genetic signature equal to that of triptolide.

### Probabilistic drug classifier

Drugs were classified as “triptolide-like” based on their dependency on PTBP1, BCL2L12, BAX, and BAK1 for causing cell death. To perform this functional genetic classification, we adapted an established approach used previously^47,48^. Briefly, for each drug-dose pair, AUC ratios between each KO and U2OS were determined. This places each drug-dose pair in a 3-dimensional functional genetic space. The “neighborhood” of transcriptional inhibitors was calculated as the mean of pairwise Euclidean distances between validated Pol II degraders (triptolide and ⍺-amanitin) in this space. The linkage ratio describes the similarity of a query compound to triptolide and ⍺-amanitin and defines how much the neighborhood expands or shrinks after the query compound is included in it. A set of negative controls, defined as a set of drugs with known mechanism of action distinct from transcriptional inhibition, is then forced to be included in the transcriptional inhibitor neighborhood, iteratively, causing the neighborhood to expand or shrink. BH3 mimetics (ABT-199, ABT-263, ABT-737), Topoisomerase inhibitors (Etoposide, Camptothecin, Idarubicin, Topotecan, and Teniposide), and ER-stress inducers (Thapsigargin, Tunicamycin, and Brefeldin A) were used to generate the false expansion. This process of “false expansion” generates an empirically defined null distribution of linkage ratios. A linkage ratio for each drug-dose pair is then determined and compared to the null distribution to derive a *P* value. *P* values are then FDR corrected using the Benjamini–Hochberg procedure.

### Data analysis and statistics

Unless otherwise noted, data analysis and statistics was performed in MATLAB (version R2024b) using built-in functions. Bar plots were generated in GraphPad Prism 10. ImageStudio 3.1.4 was used to analyze western blots. Flow cytometry analysis was performed using FlowJo version 10.8.1 software.

## SUPPLEMENTARY TABLES

**Supplementary Table 1: Mechanisms of cell death activated by Pol II degrading drugs.** Raw drug response data for triptolide and a-amanitin in the presence or absence of pathway specific inhibitors of certain death pathways (related to Fig. 1e). Canonical activators of ferroptosis, cuproptosis, parthanatos, necroptosis, and apoptosis are included as controls. Data are fractional viability measurements made using the FLICK assay.

**Supplementary Table 2: Quantification of Pol II-o inactivation and Pol II-a decay kinetics following transcriptional inhibition.** Quantified intensities for Pol II-o and Pol II-a following exposure to transcriptional inhibitors at varied doses. Data are related to Fig. 2.

**Supplementary Table 3: Raw sequencing counts from chemo-genetic profiling of triptolide treated U2OS cells.** Raw sgRNA-level counts table from chemo-genetic profiling of U2OS cells exposed to 1 µM triptolide. Data are related to Fig. 4a and b.

**Supplementary Table 4: Genetic dependencies of triptolide-induced cell death.** Gene-level drug-induced growth rates and death rates from chemo-genetic profile of triptolide in U2OS cells. Data are related to Fig. 4b.

**Supplementary Table 5: TIS score based classification to identify “TI-like” compounds.** Raw lethality data, TIS score, and classification linkage ratio used in the identification of new “TI-like” compounds. Data are related to Fig. 5a-d.

**Supplementary Table 6: Validation of new “TI-like” compounds.** Quantified intensities for Pol II-o and Pol II-a following exposure to compounds characterized as “TI-like” or “Not TI-like” at varied doses. Data are related to Fig. 5g.

---

**Extended Data FSSig. 1:**
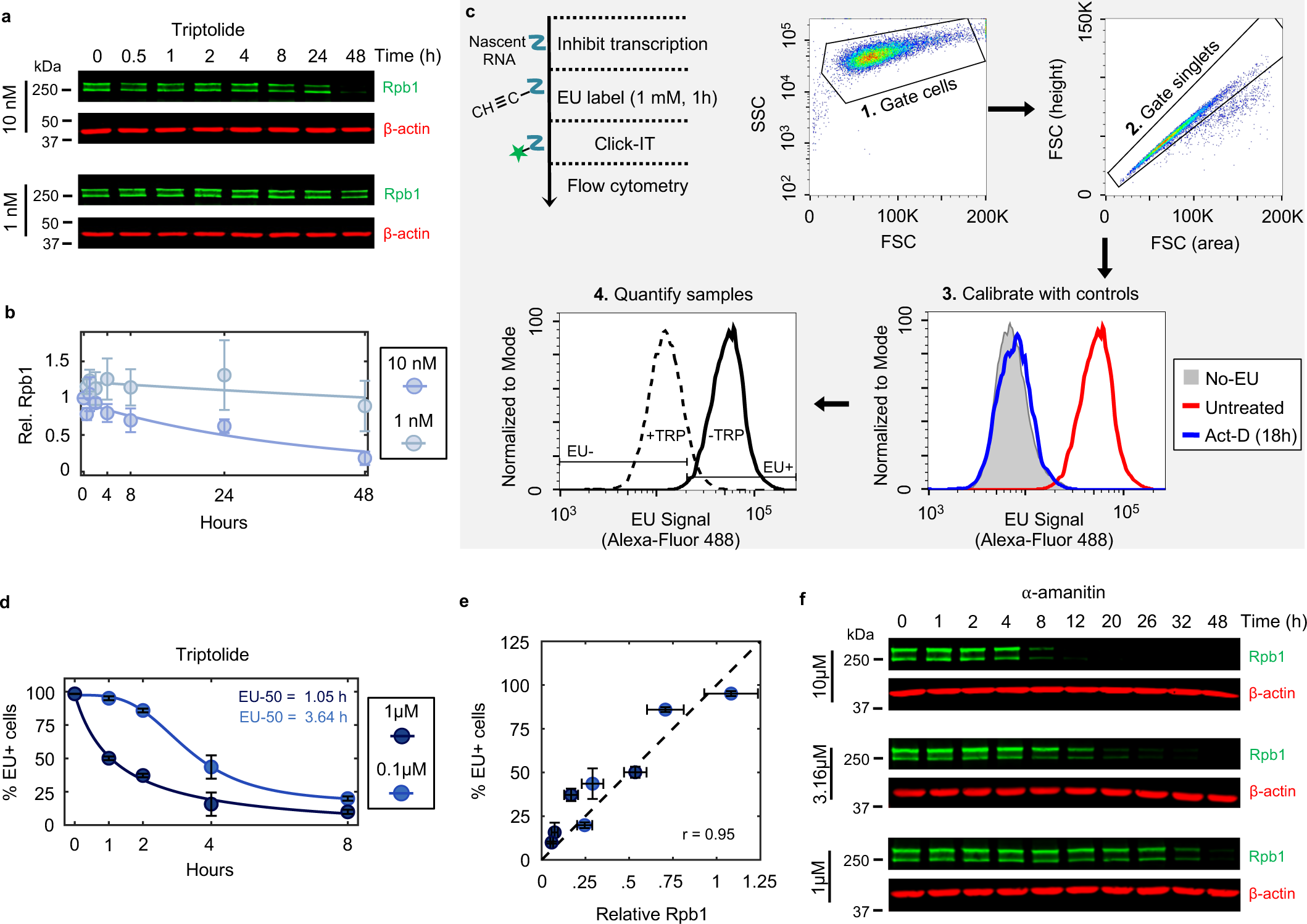
Establishing kinetics of complete transcriptional inhibition. Related to. **Fig. 1**. (**a**) Immunoblots of total Rpb1 in U2OS cells following exposure to low doses of triptolide (10 nM or 1 nM). Blots are representative of three independent biological replicates. (**b**) Quantification of Rpb1 levels shown in panel (a). Data are normalized relative to untreated levels. (**c**) Approach for measuring incorporation of 5-ethynyl uridine (EU) into newly synthesized RNAs using flow cytometry. Following labeling of nascent RNA via incubation with EU, cells are fixed, and labeled RNA is ligated to a fluorophore using click chemistry. A high dose (1 µM) and long incubation with the pan-RNA polymerase inhibitor actinomycin-D (Act-D) is used as a control to define the signal associated with no active transcription (EU-). (**d**) EU incorporation in U2OS cells before and after triptolide. Data are mean ± SD, n = 3 independent biological replicates. (**e**) Correlation between Rpb1 protein loss (quantified by immunoblot) and nascent RNA loss (quantified by EU incorporation) following exposure to 1 µM and 0.1 µM triptolide (at 1, 2, 4 and 8 hours post drug addition). Dashed line, x = y. Pearson correlation coefficient is shown. Data are mean ± SD, n = 3 independent biological replicates. (**f**) Immunoblots of total Rpb1 protein in U2OS cells following exposure to a dose range of ⍺-amanitin. Blots are representative of three independent biological replicates.

**Extended Data FSSig. 2:**
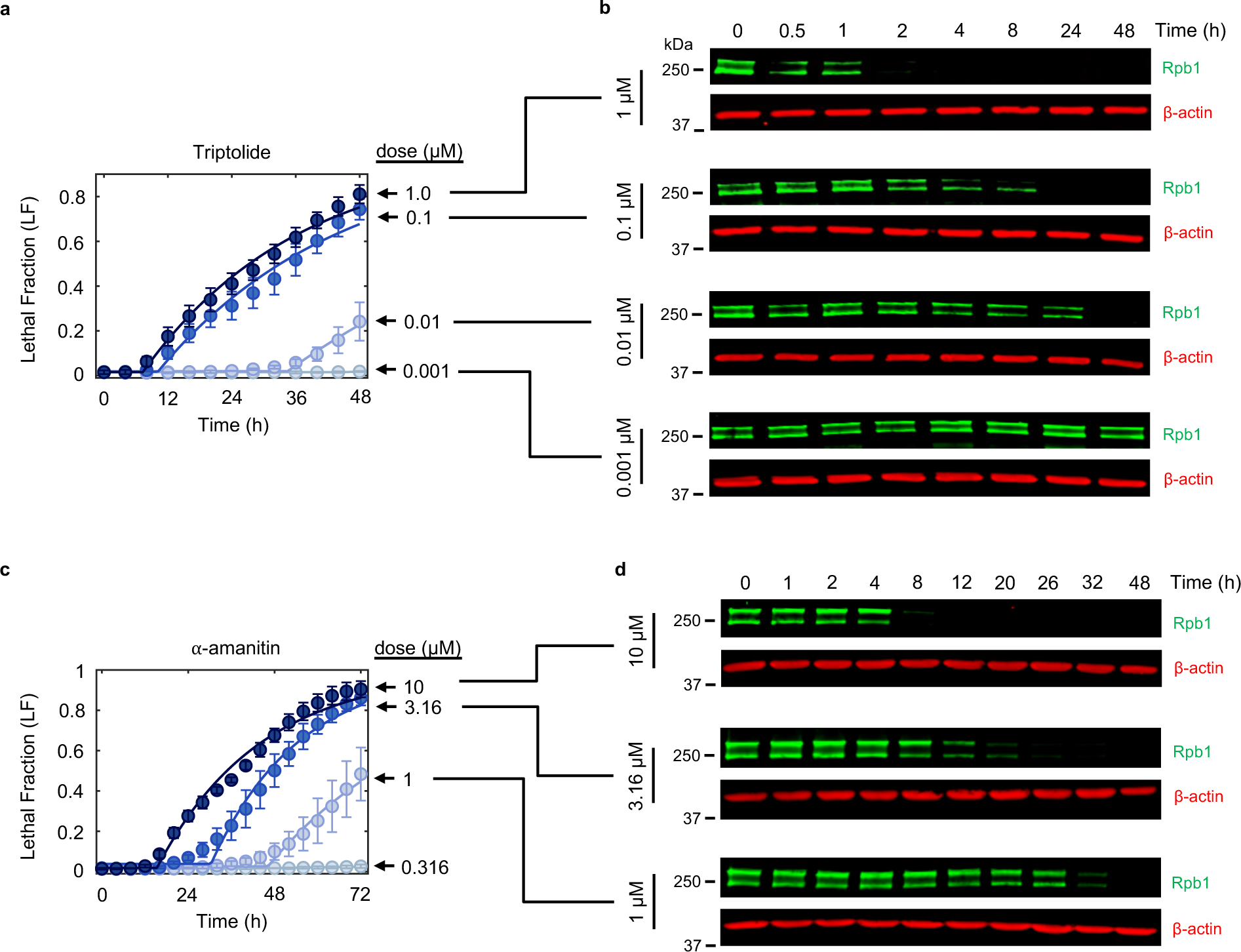
Cell death kinetics for triptolide and ⍺-amanitin across dose. Related to. **Fig. 1**. (**a**) Lethal fraction kinetics in U2OS cells for triptolide across dose, measured in STACK. Data are mean ± SD, n = 3 independent biological replicates. (**b**) Immunoblots of Rpb1 protein levels in U2OS cells following exposure to triptolide across dose. Dose associations with (a) are shown. Blots are representative of three independent biological replicates. (**c-d**) As in (a-b), but for ⍺-amanitin.

**Extended Data FSSig. 3:**
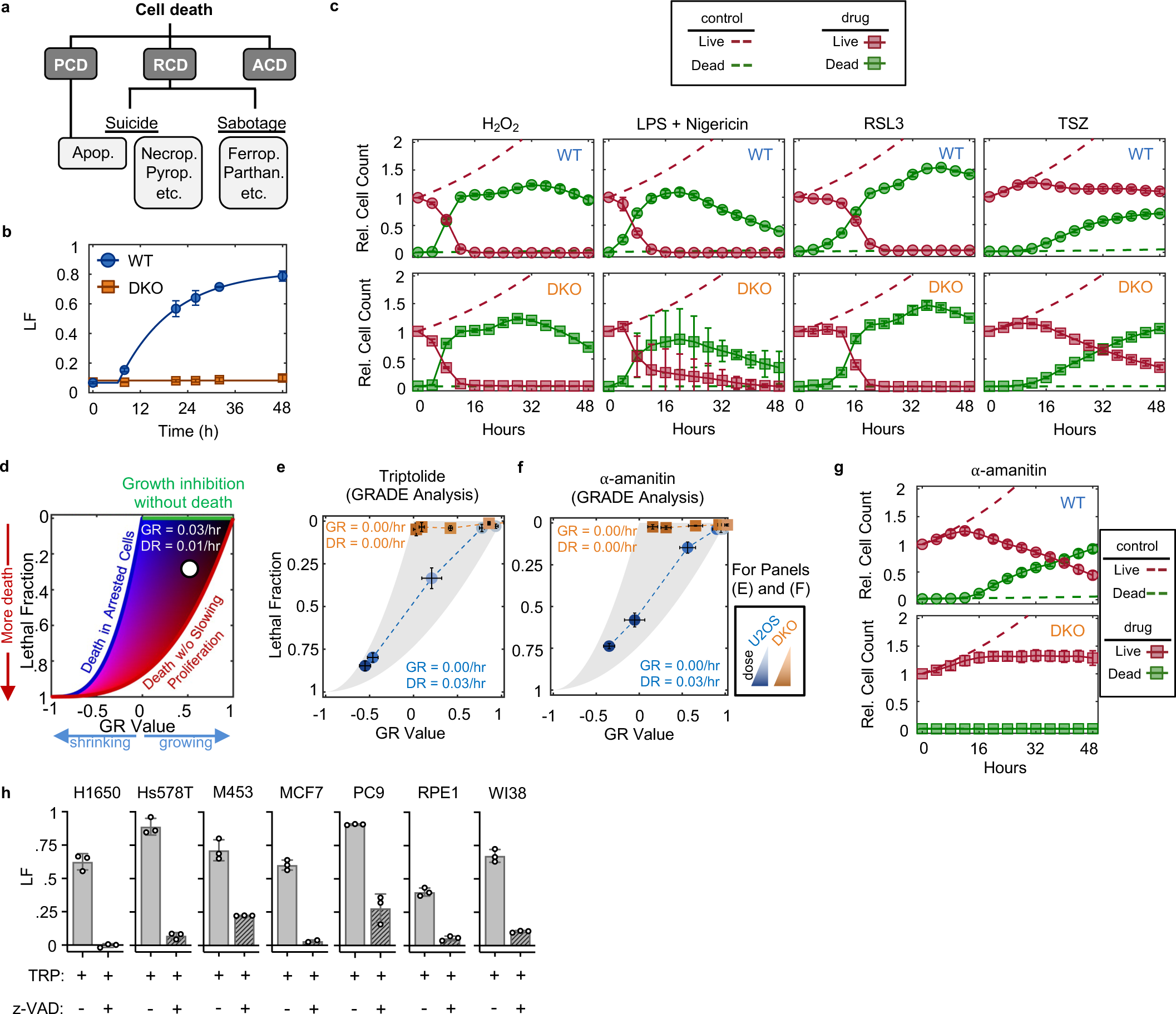
Evaluation of cell death following Pol II degradation. Related to. **Fig. 1**. (**a**) Cell death can be broadly classified into three groups: “Accidental cell death” (ACD) results from a stress that overwhelms cellular control mechanisms; “Programmed cell death” (PCD) occurs in normal development and tissue homeostasis; “Regulated cell death” (RCD) is executed by defined signaling events and effector molecules in response to exogenous stresses. RCD is further sub-divided into cell death pathways activated via signaling (cell “suicide”) or those that take advantage of cellular activities but are not necessarily due to signaling (cell “sabotage”). (**b**) LF kinetics in U2OS (WT) and U2OS^BAX-/-BAK1-/-^ (DKO) cells following exposure to the apoptotic agent ABT-737 (31.6 µM), measured in FLICK. (**c**) Live and dead cell kinetics following exposure to various non-apoptotic stimuli (Oxeiptosis, 316 µM H_2_O_2_; Pyroptosis, 0.2 µg/mL LPS + 31.6 µM Nigericin; Ferroptosis, 31.6 µM RSL3; Necroptosis, 10 ng/mL TNF⍺ + 10 µM SM-164 + 50 µM z-VAD). Data collected using the STACK assay. (**d**) Schematic of the GRADE method. Drug-induced growth and death rates can be inferred from GR/LF plots. (**e**) GRADE-based analysis of triptolide at 48 hours. Inference of growth (GR) and death (DR) rates for the highest dose (1 µM) are shown for both genotypes. (**f**) As in (e) but for α-amanitin. **(g)** Same as (c), for 10 µM ⍺-amanitin. (**h**) Lethal fraction kinetics for a panel of cell lines following 72-hour treatment with 1 µM triptolide, with- or without co-treatment with 50 µM z-VAD, measured using FLICK. (b, c, e, f, g, h) Data are mean ± SD, n = 3 independent biological replicates.

**Extended Data FSSig. 4:**
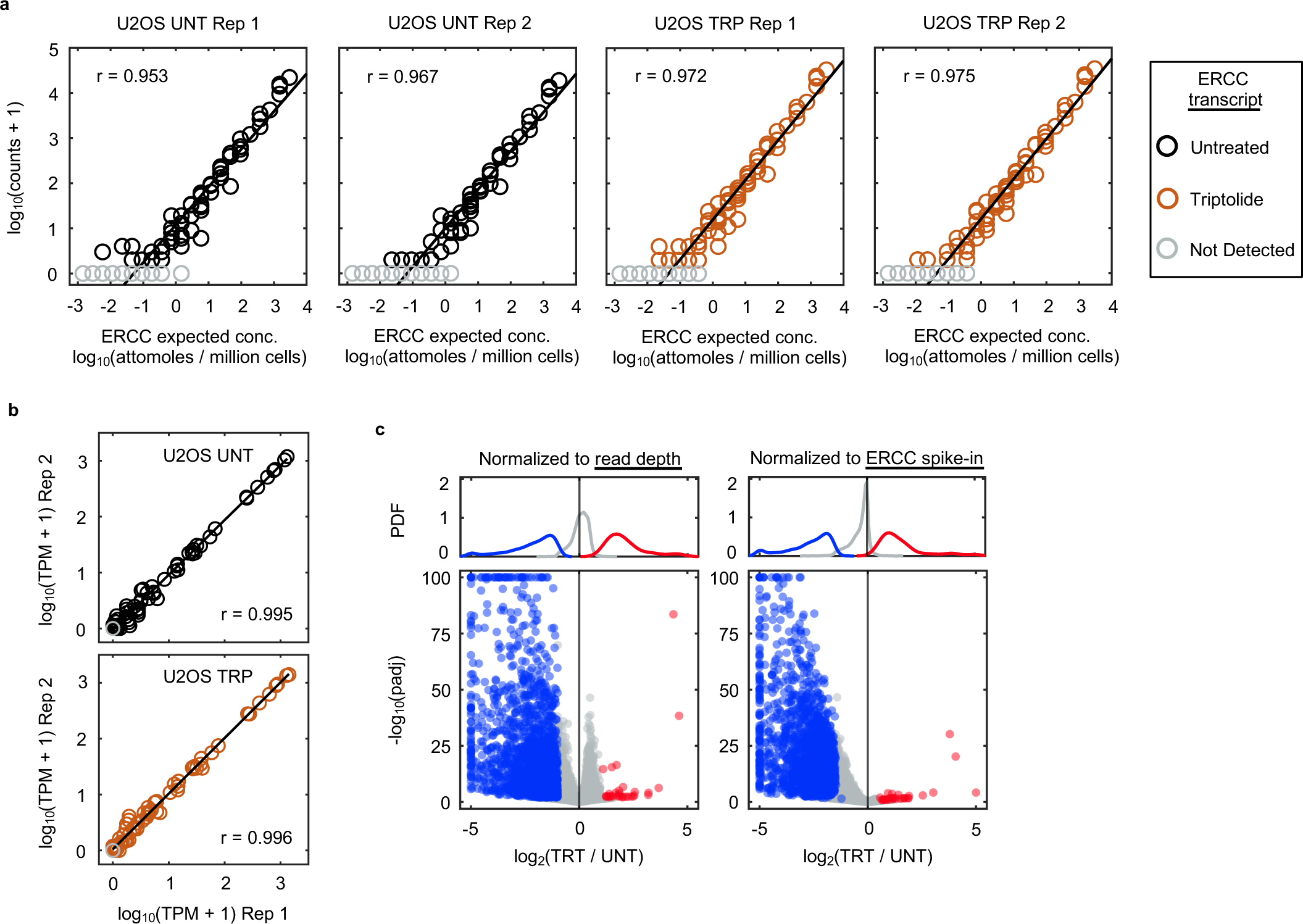
Normalization of RNAseq data to polyadenylated ERCC spike-ins. Related to. **Fig. 1**. (**a**) The expected molar amount of RNA for each of the 92 ERCC spike-in transcript standards compared to the empirically observed read counts, for replicates of U2OS RNAseq samples untreated (UNT) or treated with 1 µM triptolide (TRP) for 4 hours. Grey circles denote ERCC transcripts with zero counts. Linear regression line is shown, along with the Pearson correlation coefficient. (**b**) Transcript abundance correlation between replicates for the 92 ERCC spike-in transcript standards (TPM, transcripts per million). (**c**) Spike-in normalization is required to measure mRNA on an absolute scale. RNA-seq profiling of mRNA expression changes following exposure to 1 µM triptolide for 4 hours. (Left) mRNA expression normalized to read depth. (Right) mRNA expression normalized to ERCC spike-ins. Genes are colored based on read depth normalization: grey: no change; blue: down regulated; red: up regulated.

**Extended Data FSSig. 5:**
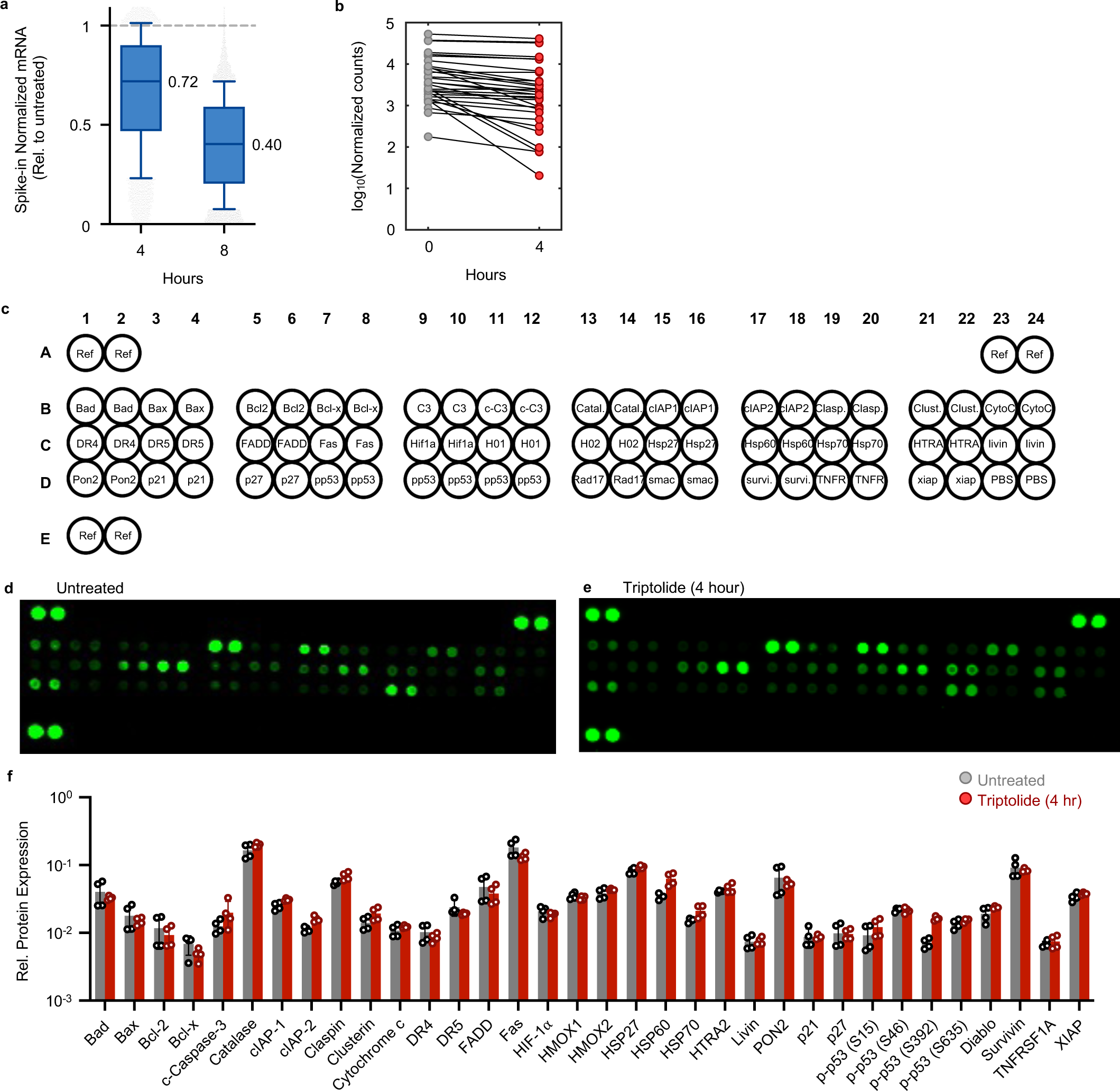
Minimal mRNA and protein decay at onset of triptolide-induced death. Related to. **Fig. 1**. (**a**) mRNA transcript abundance following 4 or 8 hours of 1 µM triptolide, measured using spike-in normalized RNAseq. Boxplots depict 10 – 90 percentile, and the median value is stated. (**b**) mRNA expression changes following exposure to 1 µM triptolide for apoptotic regulatory genes, measured using spike-in normalized RNAseq. Genes were selected to match the proteins analyzed in (c-f) in the Proteome profiler apoptosis array. (**c-f**) Proteome profiler apoptosis array. (**c**) Proteins measured in the proteome profiler array. (**d**) Untreated U2OS cells. Data are representative of two independent biological replicates. (**e**) As in (d), for U2OS cells treated with 1 µM triptolide for 4 hours. (**f**) Quantification of (d-e). Data are means ± SD of two independent biological replicates and two technical replicates, normalized to the highest expression protein (total Pro-Caspase-3). Significant differences between treated and untreated conditions for each epitope were determined using unpaired t tests with Welch correction, and the two-stage step-up method of Benjamini, Krieger and Yekutieli was used to correct for multiple comparisons. No proteins were identified as being significantly different, except for phospho-p53 (S392) (q = 0.001).

**Extended Data FSSig.6:**
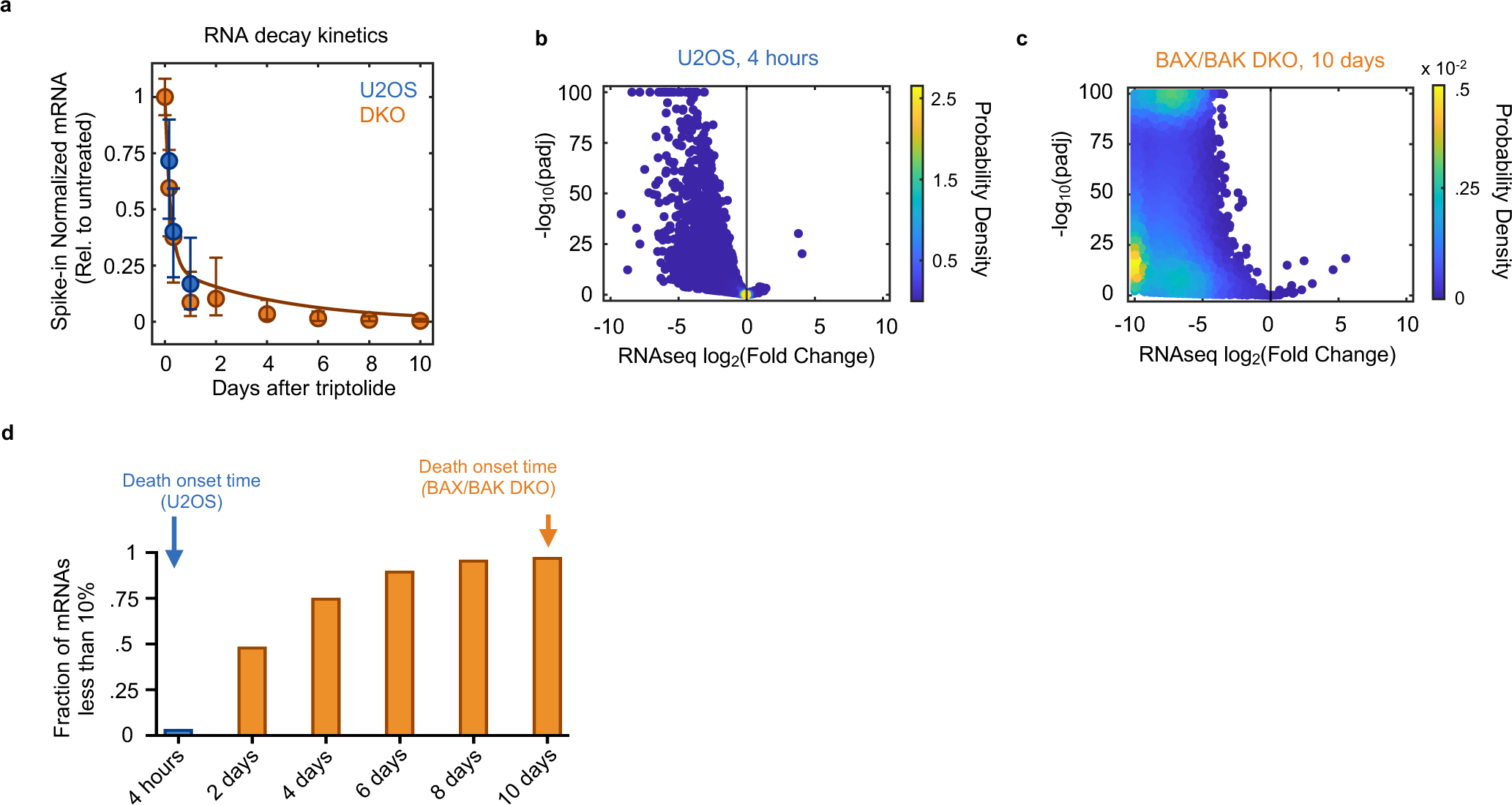
mRNA decay following long-term exposure to triptolide. Related to. **Fig. 1**. (**a**) Decay kinetics for mRNAs in U2OS and BAX/BAK DKO (DKO) after 1 µM triptolide exposure for the indicated times, measured using spike-in normalized RNAseq. Datapoints indicate median, 25^th^, and 75^th^ percentiles. (**b**) Spike-in normalized mRNA fold changes in U2OS cells following 4 hours of 1 µM triptolide. (**c**) Spike-in normalized mRNA fold changes in *BAX/BAK* DKO cells following 10 days of 1 µM triptolide. (**d**) Fraction of mRNAs that have decayed beyond 90% of untreated steady-state abundance. U2OS (blue) and *BAX/BAK* DKO cells (orange) are shown, and their respective death onset times are noted.

**Extended Data FSSig. 7:**
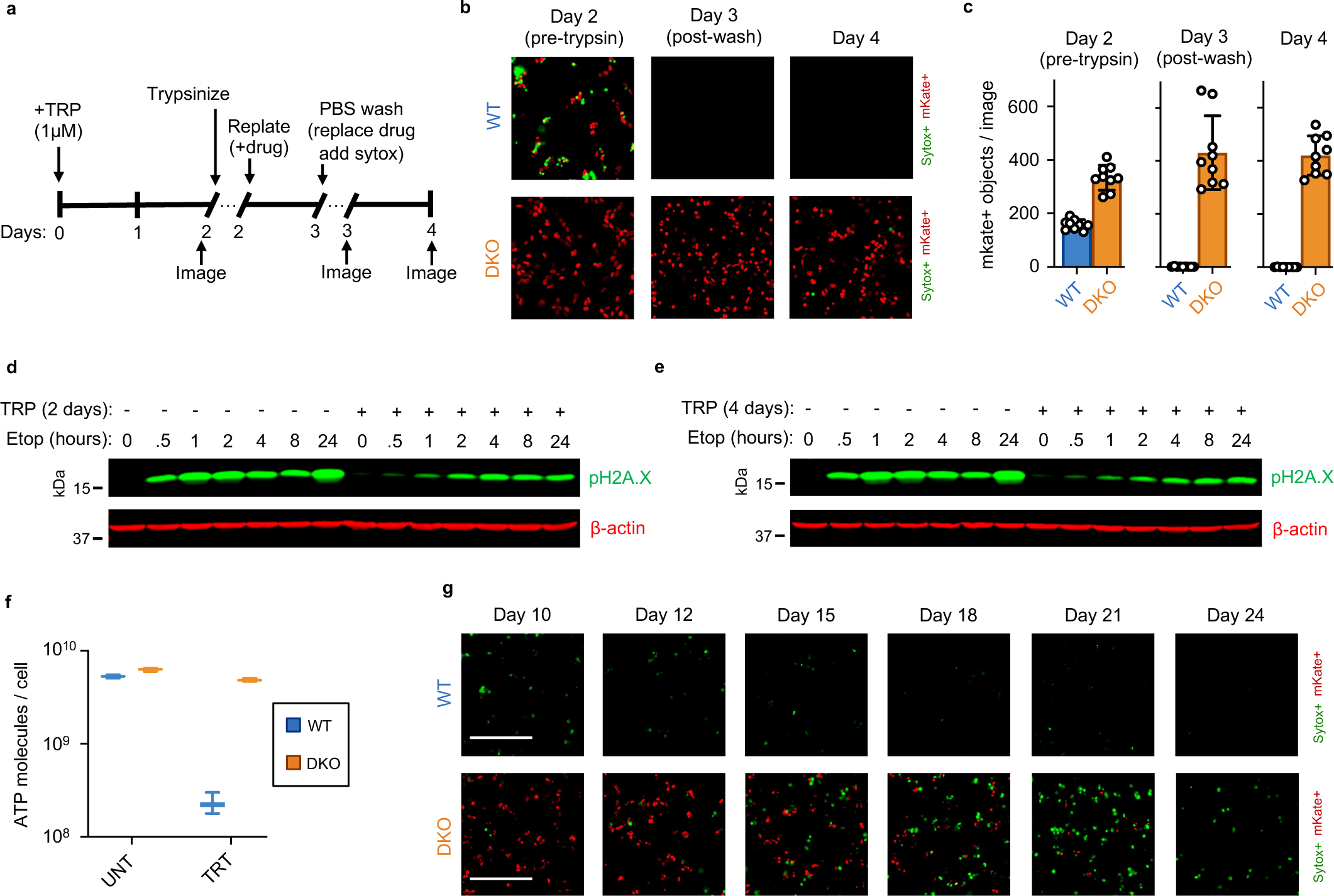
Viability of *BAX*/*BAK* DKO cells during long-term exposure to triptolide. Related to. **Fig. 1** (**a**) Experimental overview for testing the ability of apoptotic deficient cells to reattach to plastic following trypsinization. (**b**) Representative images of triptolide treated cells before and after trypsinization and subsequent replating. (**c**) Quantification of mKate+ nuclei per image before and after trypsinization and subsequent replating, associated with (b). (**d**) Immunoblot depicting intact DNA damage response signaling in both untreated and triptolide-treated BAX/BAK DKO cells. Cells were first treated with or without 1 µM triptolide for 2 days to induce Pol II degradation, following by addition of 31.6 µM Etoposide to induce DNA damage. pH2A.X (S139) levels were monitored over time following Etoposide. (**e**) As in (d), except BAX/BAK DKO cells were exposed to triptolide for 4 days. (**f**) ATP molecules per cell following a 4-day treatment with 1 µM triptolide in U2OS or BAX/BAK DKO cells. ATP was measured using Cell Titer Glo, and cell number was normalized using FLICK. Mean, minimum, and maximum values of n = 60 independent biological replicates are shown. (**g**) Long-term survival of BAX/BAK DKO cells following 1 µM triptolide (associated with Fig. 1f). Representative images for three independent biological replicates is shown.

**Extended Data FSSig. 8:**
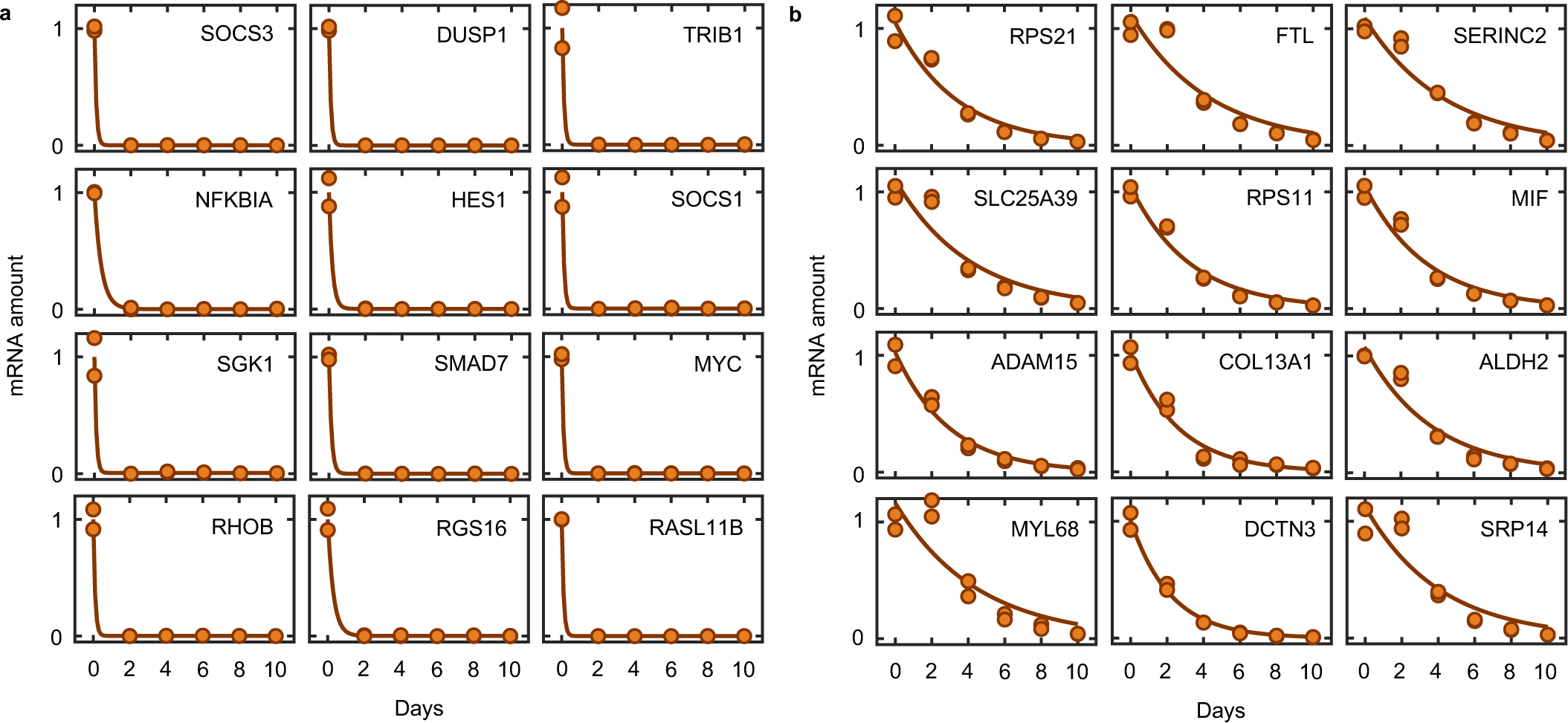
Decay kinetics of short and long-lived transcripts. Related to. **Fig. 1**. (**a**) mRNA decay kinetics in DKO cells following long-term exposure to 1 µM triptolide, measured using spike-in normalized RNAseq. 12 shortest half-life transcripts, defined from a meta-analysis of half-life measurements Agarwal and Kelley (2022). (**b**) As in (a), for the 12 longest half-life transcripts.

**Extended Data FSSig. 9:**
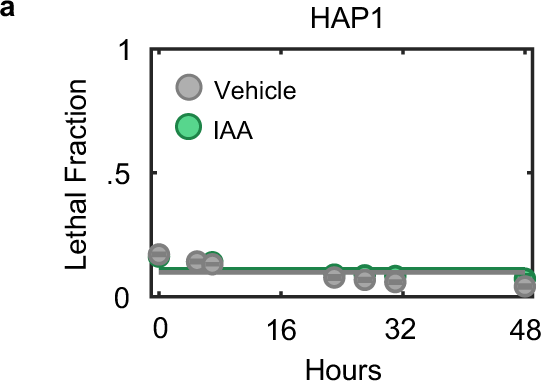
Cell death in HAP1-RPB1-AID cells is not caused by auxin itself. Related to. **Fig. 1** (**a**) Lethal fraction kinetics in parental HAP1 cells following exposure to 500 µg/mL auxin (IAA, 3-indoleacetic acid) or vehicle, measured in FLICK. Data are mean ± SD, n = 3 independent biological replicates.

**Extended Data FSSig. 10:**
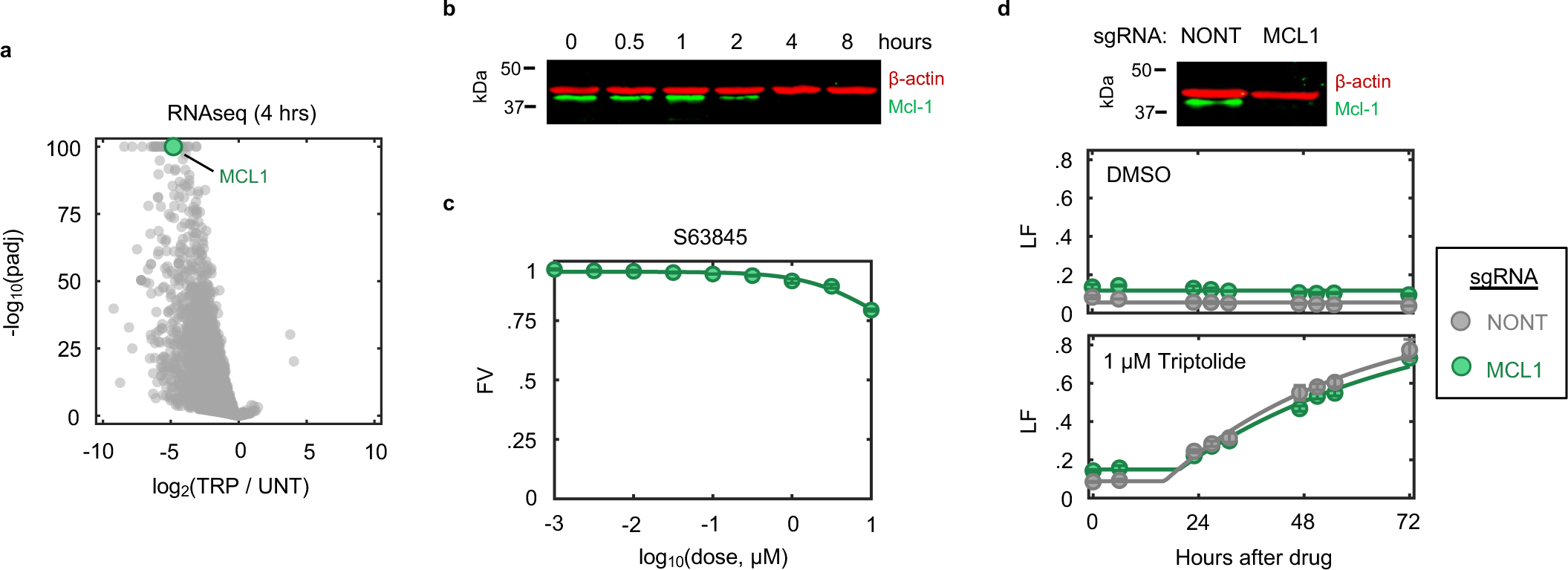
Loss of Mcl-1 protein following transcriptional inhibition is not causal for apoptosis. Related to. **Fig. 2**. (**a**) mRNA expression following 4-hour exposure to 1 µM triptolide in U2OS cells, highlighting the rapid loss of MCL1. (**b**) Immunoblot of Mcl-1 protein levels in U2OS cells following exposure to 1 µM triptolide. (**c**) U2OS cells are insensitive to the Mcl-1 inhibitor S63845. Fractional viability (FV) measured after 72 hours of drug exposure, measured in FLICK. Data are mean ± SD for n = 3 independent biological replicates. (**d**) Knockout of MCL1 is not lethal in U2OS cells and does not impact the lethality induced by triptolide. (Top) Immunoblot of Mcl-1 protein levels in U2OS cells expressing sgRNA targeting MCL1 or nontargeting sgRNA. (Bottom) Lethal fraction (LF) kinetics between the two genotypes following exposure to triptolide or vehicle (DMSO, 0.1%). LF kinetics were measured in FLICK, and data are mean ± SD for n = 3 independent biological replicates. Protein lysates used for the immunoblot were taken from the same pool of cells used for LF measurements and were collected at the time of drugging (“0 hours”).

**Extended Data FSSig. 11:**
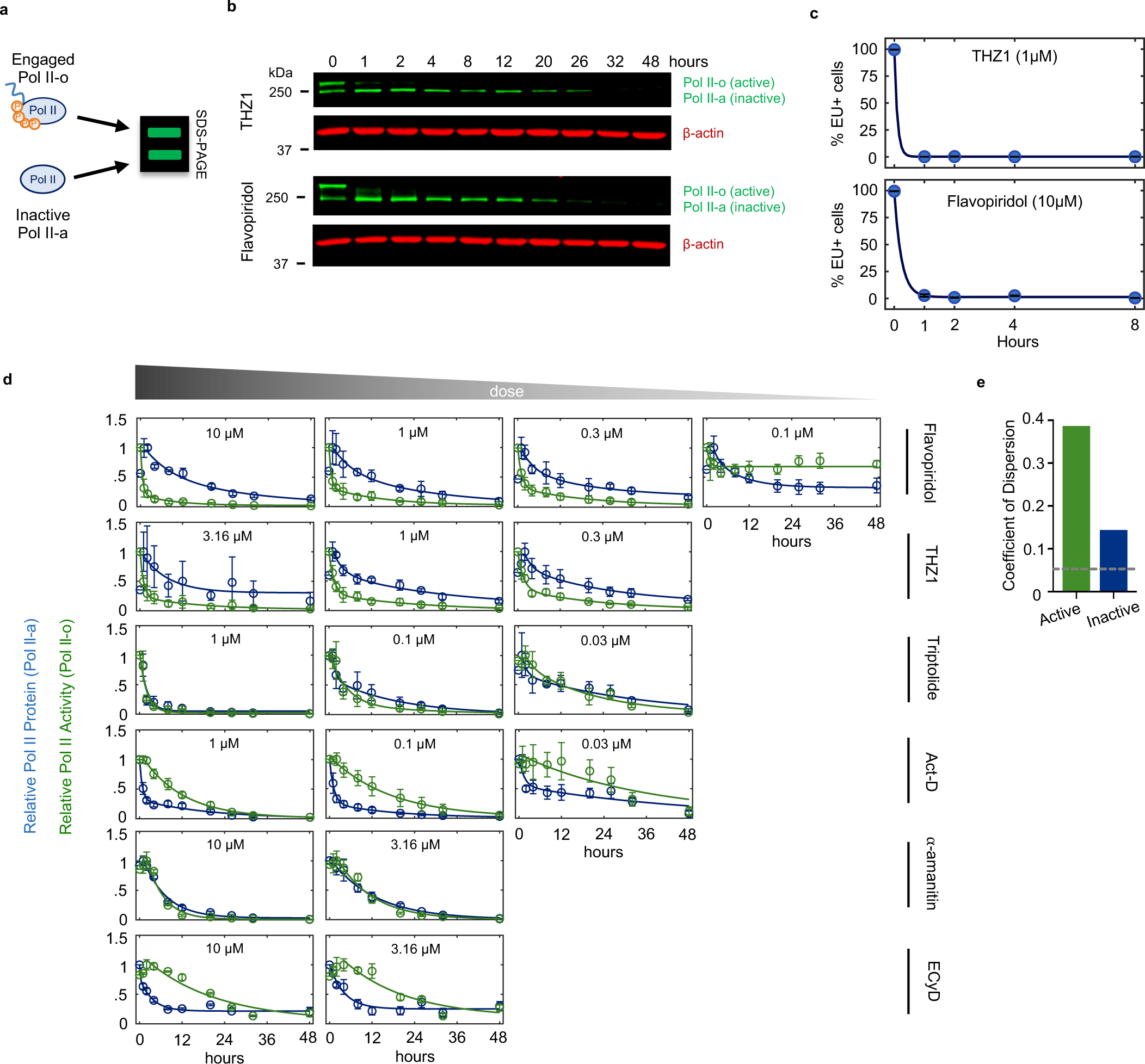
Variation in kinetics of Pol II inhibition across transcriptional inhibitors. Related to. **Fig. 2**. (**a**) Pol II protein primarily exists in two forms. Pol II-o is hyper-phosphorylated along the carboxy-terminal domain (CTD) of the protein and is the actively elongating pool of Pol II. The hypo-phosphorylated form, called Pol II-a, represents all other inactive forms of Pol II, including preinitiation complex (PIC) bound, promoter paused, early pause released, and free Pol II. Pol II-o and Pol ll-a can be distinguished by a band shift on a gel. (**b**) Example immunoblot of Rpb1 levels following exposure to two fast acting transcriptional inhibitors, THZ1 (1 µM), and Flavopiridol (10 µM). These drugs rapidly decrease Pol II-o levels. (**c**) EU incorporation into nascent RNA following fast acting inhibitors shown in (b), validating that loss of Pol II-o levels is functionally equivalent to loss of transcription. Data are mean ± SD for n =3 independent biological replicates. (**d**) Quantification of Pol II-o (green) and Pol II-a (blue) protein levels across different lethal doses of transcriptional inhibitors. Data were collected using quantitative immunoblotting and fit to a two-phase exponential decay function. Act-D, Actinomycin-D; ECyD, Ethynylcytidine. Data are mean ± SD for n =3 independent biological replicates. (**e**) Coefficient of Dispersion (CoD) of LF_50_ times for cell death kinetics aligned to active or inactive Pol II loss, as shown in Fig. 2c,e. Gray dashed line denotes the CoD for the “perfect” alignment, defined by arbitrarily shifting each kinetic curve such that maximum alignment is achieved, implemented using the MATLAB function alignsignals with method “risetime”.

**Extended Data FSSig. 12:**
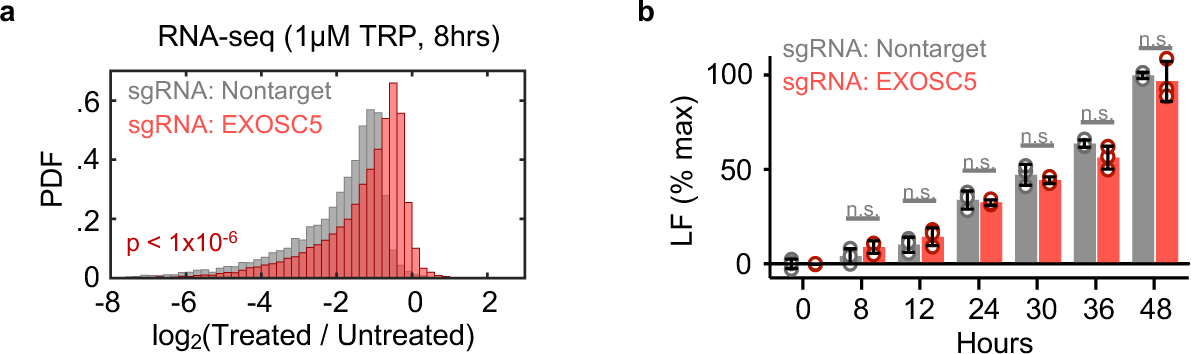
Effects of triptolide in EXOSC5 knockout cells. Related to. **Fig. 3**. (**a**) Histogram of spike-in normalized absolute mRNA fold changes following 8-hr exposure to 1 µM triptolide (TRP) in U2OS cells expressing sgRNA targeting EXOSC5 or nontargeting sgRNA. Two-sided KS test *p* value is shown. (**b**) Comparison of lethal fraction (LF) levels between cell types in (a) at various time points following exposure to 1 µM TRP, measured using the FLICK assay. Data normalized to max LF in cells expressing nontargeting sgRNA. Data are mean ± SD for n = 3 independent biological replicates. Wilcoxon rank sum *p* value shown (n.s. > 0.05).

**Extended Data FSSig. 13:**
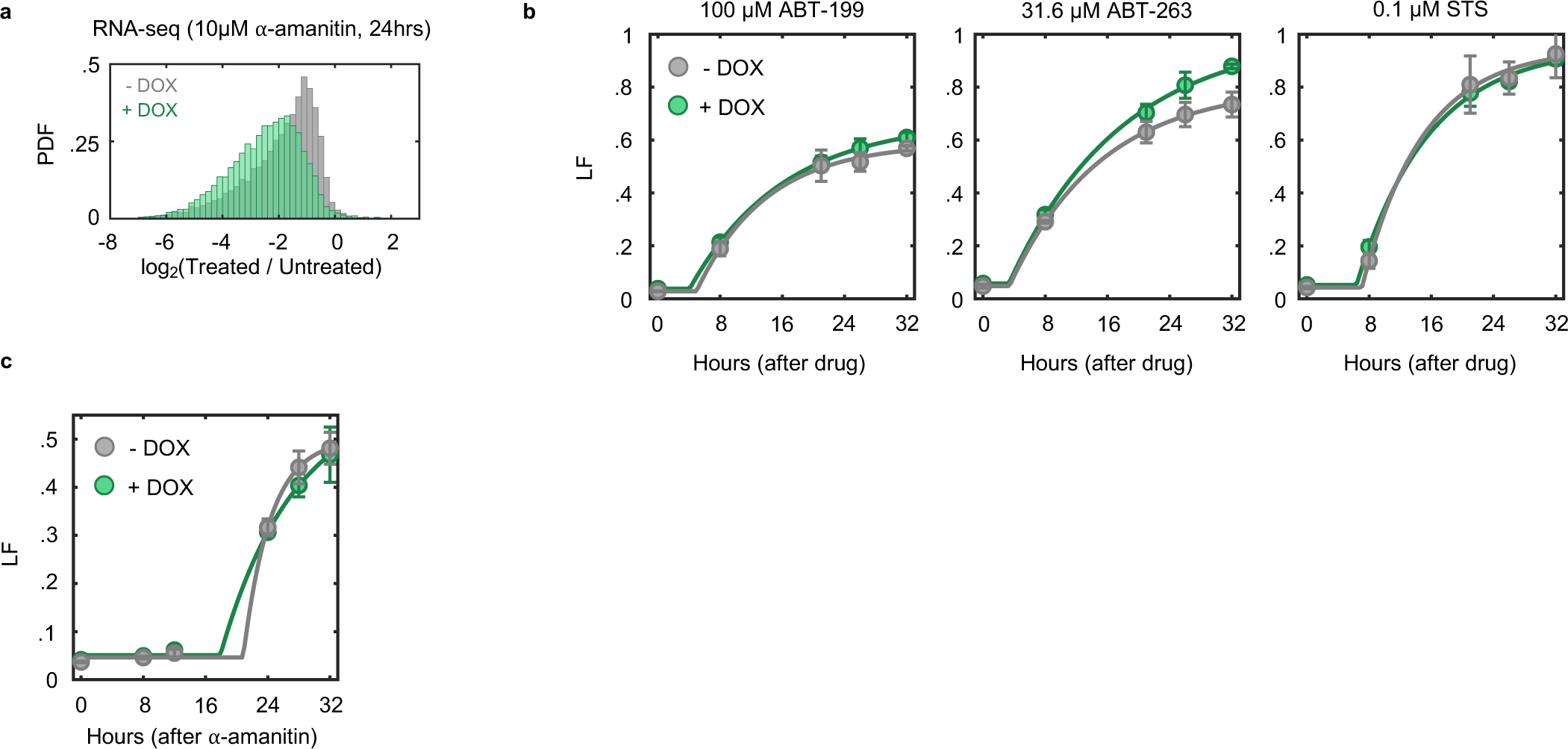
Characterization of the Pol II switchover system. Related to. **Fig. 3**. (**a**) Histogram of spike-in normalized absolute mRNA fold changes following 24-hr exposure to 10 µM ⍺-amanitin. (**b**) Lethal Fraction (LF) kinetics following exposure to general apoptotic agents in Pol II switchover cells with- or without doxycycline (DOX)-induced expression of RPB1-N792D-ΔCTD. LF measured using the FLICK. STS = Staurosporine. (**c**) LF kinetics in cells with- or without expression of a ⍺-amanitin sensitive *RPB1-*ΔCTD transgene, treated with 10 µM ⍺-amanitin and measured using FLICK. For (b-c), mean ± SD shown, n = 3 independent biological replicates.

**Extended Data FSSig. 14:**
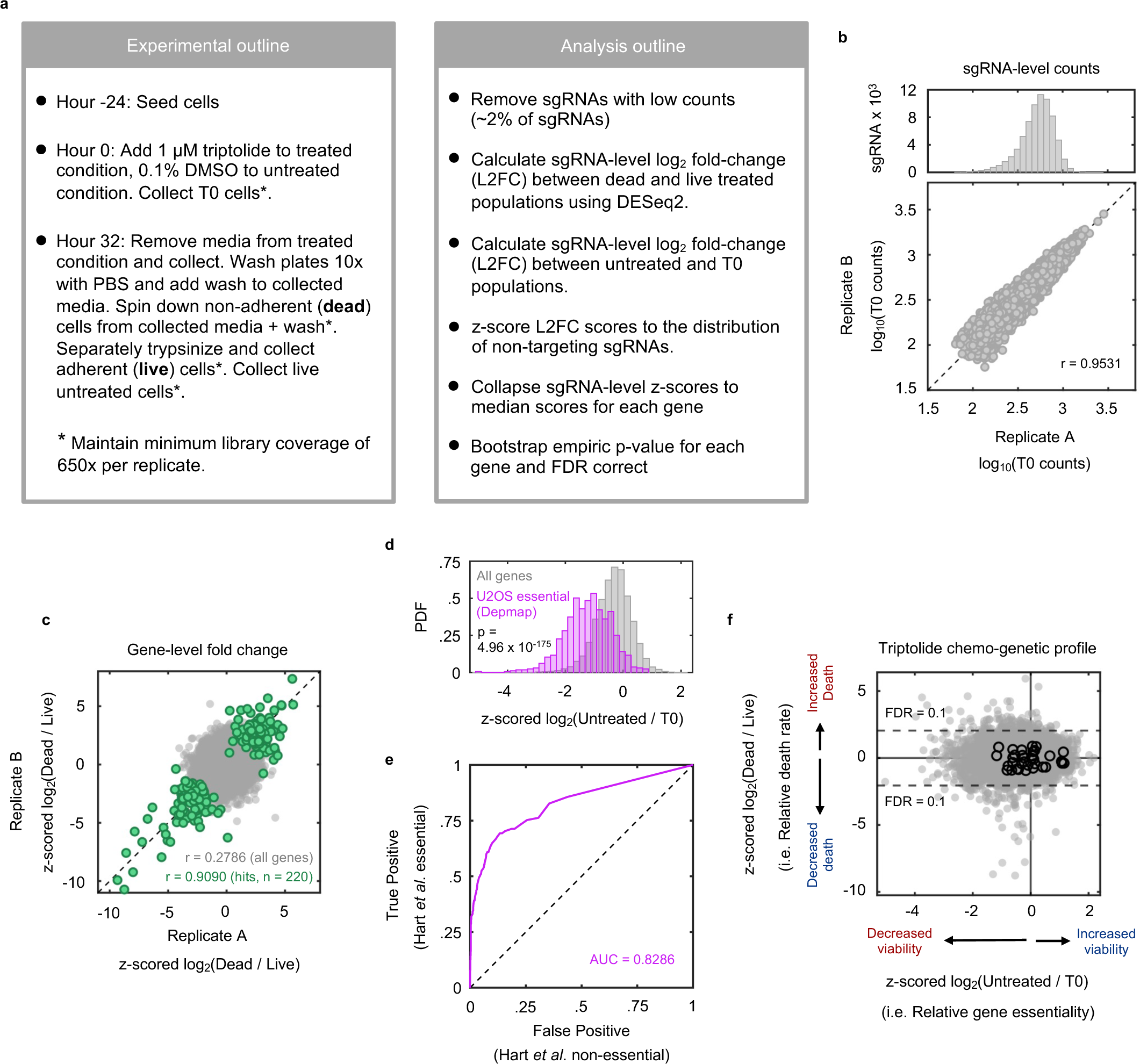
Chemo-genetic profiling strategy and quality assessment. Related to. **Fig. 4**. (**a**) Overview of experimental and analytical process for the chemo-genetic screen. (**b**) (Top) Representative example sgRNA count distribution for replicate A of the T0 condition. (Bottom) Representative example correlation of sgRNA counts between two replicates of the same condition. Dashed line, x = y. Pearson correlation coefficient is shown. (**c**) Correlation of gene-level L2FC scores between two replicates treated with triptolide. (**d**) Distribution of gene-level L2FC scores for essential genes versus all genes in untreated vs T0 comparison. Two-sided KS test *p* value is shown. (**e**) ROC curve depicting the sensitivity and specificity of the untreated versus T0 screening comparison to classify previously established essential and non-essential genes. (**f**) Conceptual overview of metrics comprising the chemo-genetic profile of triptolide. The x-axis describes the effect a gene knockout has on the viability of a cell in untreated conditions. The y-axis describes the effect a gene knockout has on the cell death rate in the context of triptolide. Scores for all genes (gray, filled), and non-targets (black, empty) are shown.

**Extended Data FSSig. 15:**
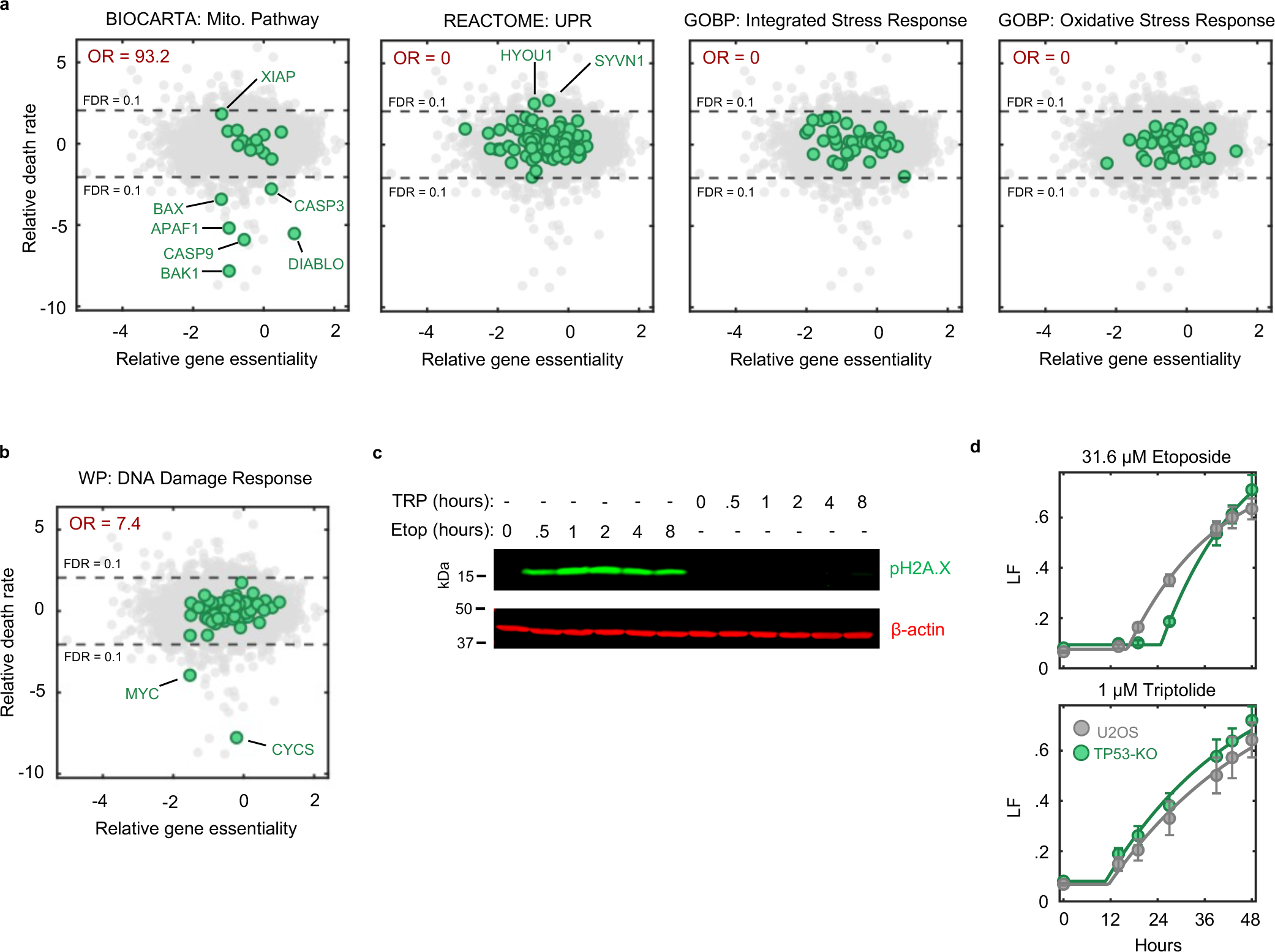
Triptolide-induced apoptosis does not depend on known stress response pathways. Related to. **Fig. 4**. (**a**) Gene-level chemo-genetic profiling data for U2OS cells treated with 1 µM triptolide, highlighting genetic signatures associated with stress response pathways. Signatures defined from the Molecular Signatures Database (MSigDB). The mitochondrial pathway, involved in intrinsic apoptosis, are enriched (far left) among the genes whose knockout decreases triptolide-induced death. The unfolded protein response (UPR), integrated stress response (ISR), and the oxidative stress response are not enriched. Odds ratio for a one-tailed Fisher’s exact test is shown (associated adjusted p-values are shown in Fig. 5D). (**b**) As in (a), but for genes associated with the DNA Damage Response (DDR). Two hits, MYC and CYCS, are more generally involved in apoptosis and are not specific to the DDR. (**c**) Immunoblot of phosphorylated H2A.X (Ser139) levels following 31.6 µM etoposide or 1 µM triptolide exposure in U2OS cells, demonstrating that triptolide does not induce DDR signaling, as compared to the DNA damaging drug etoposide. (**d**) Lethal fraction (LF) kinetics following exposure to etoposide or triptolide in U2OS cells or clonal U2OS-TP53 knockout cells, measured in FLICK. Loss of TP53 delays cell death following DNA damage, as previously described (Honeywell, et al., 2024). Loss of TP53 has no effect on cell death following triptolide. Data are mean ± SD for n = 9 independent biological replicates.

**Extended Data FSSig. 16:**
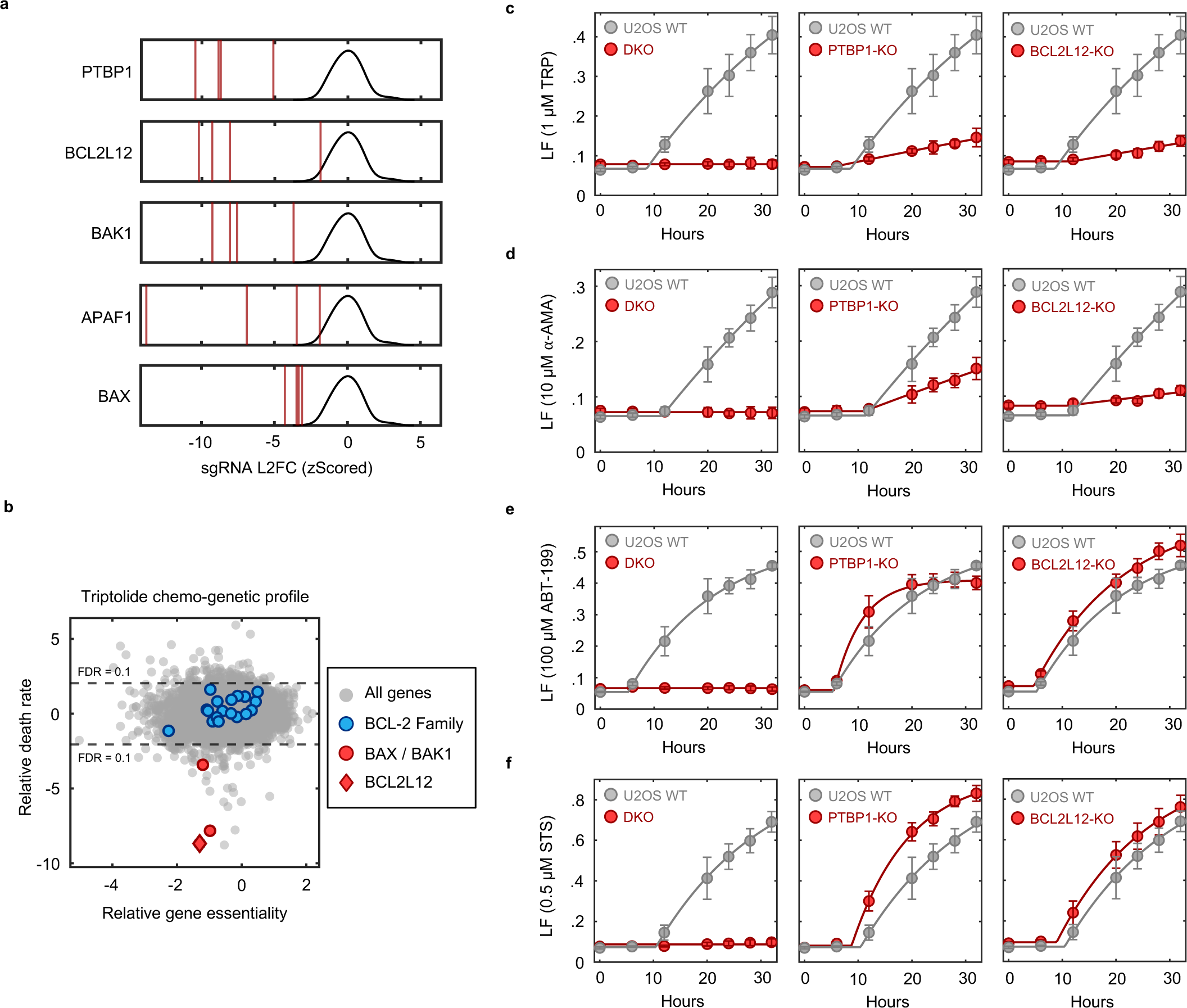
Validation of PTBP1 and BCL2L12 as unique genetic dependencies of lethality following Pol II degradation. Related to. **Fig. 4**. (**a**) sgRNA-level log2 fold changes (dead population / live population) for PTBP1 and BCL2L12 in the context of 1 µM triptolide. BAK1, APAF1, and BAX are core apoptotic regulators and act as controls. Fold changes for 4 sgRNAs per gene (red vertical lines) were z-scored to the distribution of nontargeting guides (black). (**b**) Gene-level chemo-genetic profiling data for U2OS cells treated with 1 µM triptolide. BCL-2 family proteins regulate BAX and BAK1 activity. BCL2L12 is the only BCL-2 family protein identified that modulates cell death following triptolide (apart from BAX and BAK1 themselves). Highlighted BCL-2 family members in blue are MCL1, BCL2L1, BCL2L14, BCL2L10, BCL2A1, BCL2L2, BCL2L13, BAD, BOK, BMF, BCL2, BIK, BCL2L15, BCL2L11, and BID. (**c-f**) Cell death kinetics measured using FLICK in U2OS wild-type cells compared to BAX/BAK DKO (left), PTBP1-KO (middle), and BCL2L12-KO cells (right). **(c)** 1 µM triptolide. **(d)** 10 µM ⍺-amanitin. **(e)** 100 µM ABT-199. **(f)** 0.5 µM Staurosporine. Data are mean ± SD for n = 6 independent biological replicates. Panels (c) and (d) are Pol II-degrading transcriptional inhibitors. Panels (e) and (f) are canonical apoptotic activators.

**Extended Data FSSig. 17:**
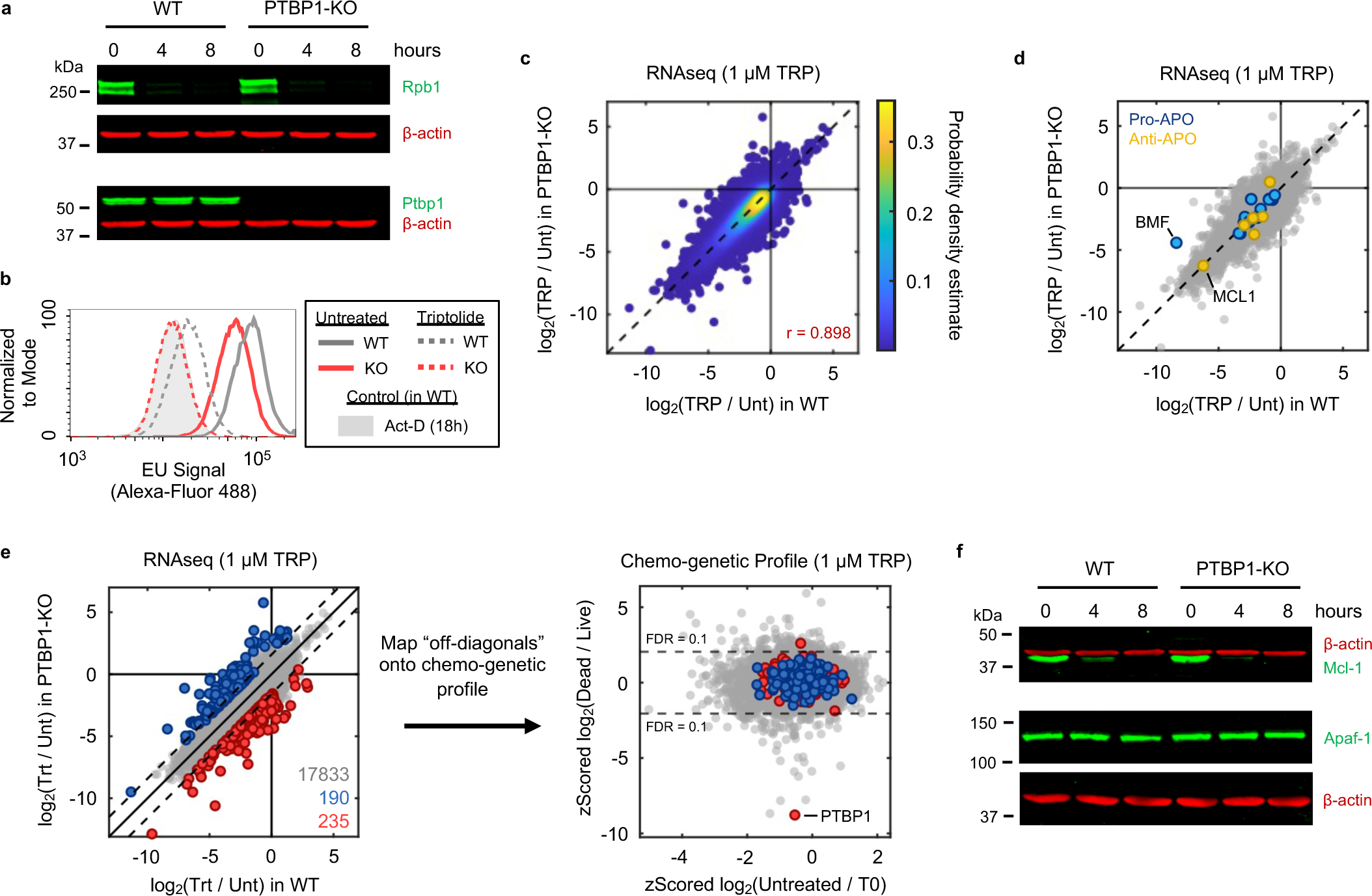
PTBP1 knockout has no effect on Pol II degradation, loss of nascent transcription, or loss of mRNA, following triptolide exposure. Related to. **Fig. 4**. (**a**) Immunoblots for Rpb1 protein levels following exposure to 1 µM triptolide in U2OS and U2OS-PTBP1-KO. (**b**) EU incorporation into nascent RNA in U2OS and U2OS-PTBP1-KO. Cells treated with or without 1 µM triptolide for 8 hours prior to EU labeling. Populations are representative of 3 biological replicates. (**c**) RNAseq comparison of drug-induced transcriptional changes in wild-type (x-axis) and PTBP1-KO (y-axis) cells, measured 8 hours following treatment with 1 µM triptolide. Data were normalized using ERCC spike-ins. Dashed line, x = y. Correlation coefficient shown is for all genes. (**d**) RNAseq data as in (c), highlighting that established anti-apoptotic (Anti-APO) and pro-apoptotic (Pro-APO) regulators show similar expression changes following triptolide in WT and PTBP1-KO backgrounds. Anti-APO: BCL2, MCL1, BCL2L10, BCL2A1, BCL2L2, XIAP. Pro-APO: BCL2L14, BCL2L13, BAD, BOK, BMF, BIK, BCL2L11, BID, BAK1, BAX, APAF1, CASP3, CASP7, CASP8, CASP9, CYCS, DIABLO. (**e**) (Left) RNAseq data as in (c), identifying rare outlier genes whose expression is decreased more (red) or less (blue) in PTBP1-KO cells compared to WT cells following triptolide. The threshold for an “outlier” is depicted by the dashed lines, denoting 2.5 standard deviations from the identity line (x = y). Number of outliers shown. (Right) Mapping of expression outliers onto the chemo-genetic profile for triptolide. (**f**) Immunoblots for Mcl-1 and Apaf-1 protein levels following exposure to 1 µM triptolide in wild-type U2OS cells and U2OS-PTBP1-KO clones.

**Extended Data FSSig. 18:**
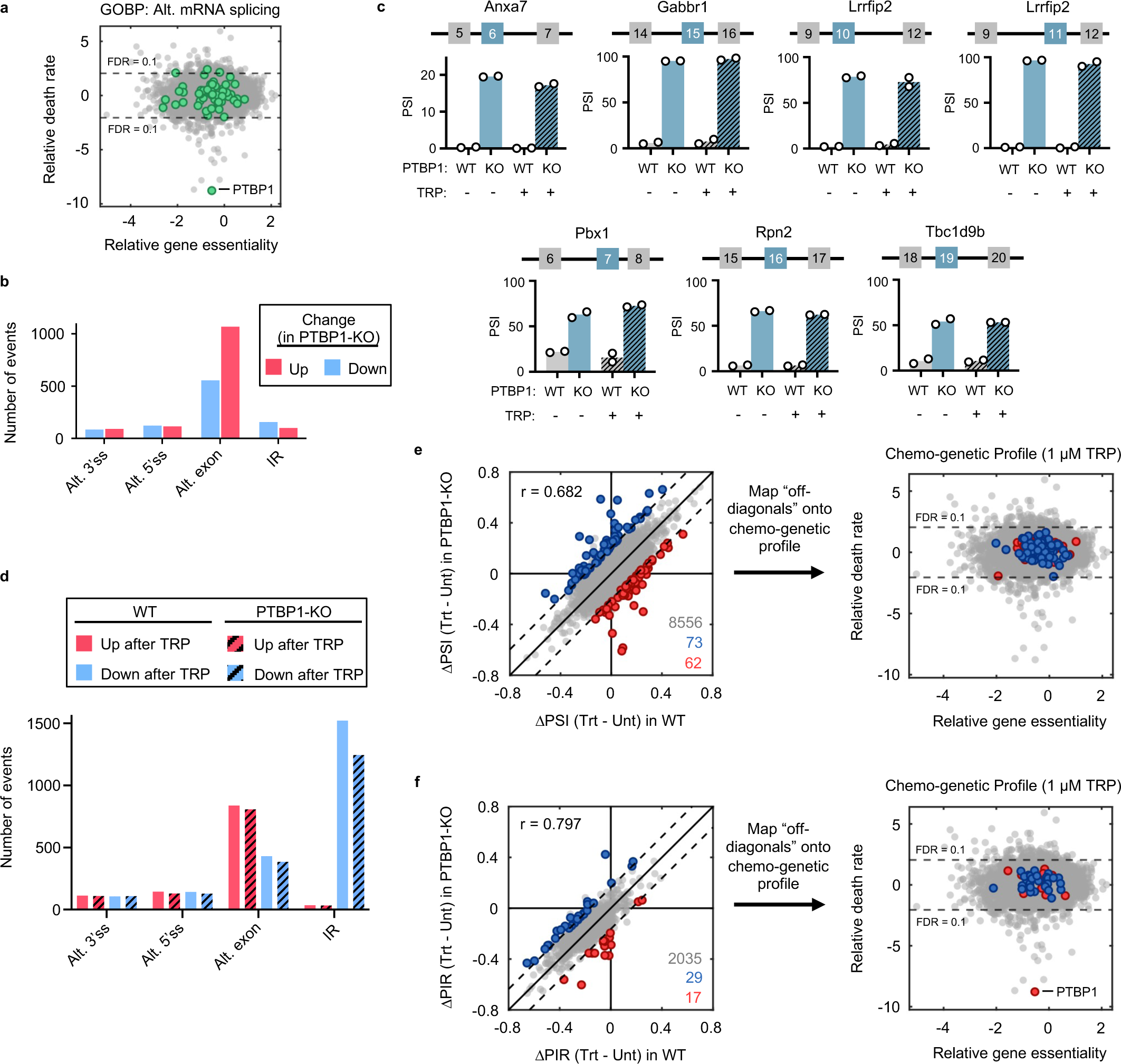
The role of PTBP1 in triptolide-induced lethality is unrelated to alternative splicing. Related to. **Fig. 4**. (**a**) Chemo-genetic profiling data for U2OS cells treated with 1 µM triptolide highlighting genes that regulate alternative splicing, as defined by MSigDB. (**b**) Number and type of significant PTBP1-dependent splicing events identified by comparing U2OS and U2OS-PTBP1-KO RNAseq datasets. (**c**) Previously annotated PTBP1-dependent exon exclusion events. PSI: percent spliced in for the blue colored exon. Data are with- or without 1 µM triptolide for 8 hours. (**d**) Significant splicing alterations induced following 8-hour exposure to 1 µM triptolide in either U2OS (solid bars) or U2OS-PTBP1-KO (dashed bars) cells. (**e**) (Left) Comparison of drug-induced splicing changes in U2OS (x-axis) and U2OS-PTBP1-KO (y-axis) cells, measured 8 hours following treatment with 1 µM triptolide. Data include alternative 3’ splice site, alternative 5’ splice site, and alternative exon usage. Dashed lines denote 2.5 standard deviations from the identity line (x = y), and off-diagonals are highlighted. (Right) Mapping of outliers onto the chemo-genetic profile for triptolide. (**f**) As in (e), but specifically for intron retention (IR).

**Extended Data FSSig. 19:**
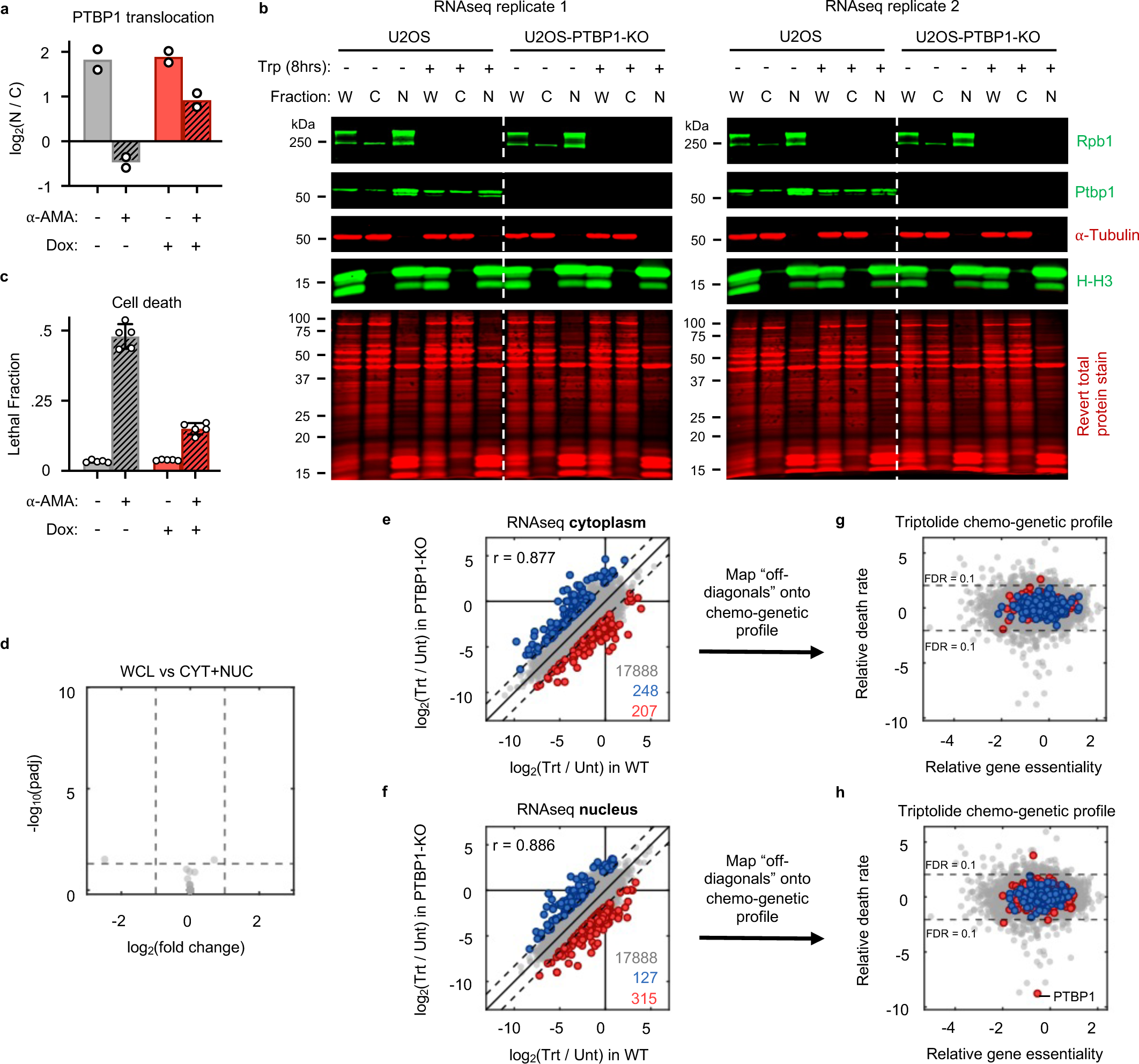
Evaluation of PTBP1 nuclear export following Pol II degradation. Related to. **Fig. 4**. (**a**) Quantification of the nuclear-to-cytoplasmic ratio of PTBP1 protein before and after exposure to 10 µM ⍺-amanitin for 12 hours in Pol II switchover cells, measured using immunoblotting. (**b**) PTBP1 translocation following triptolide (Trp) exposure. These samples were also used for RNA sequencing of nuclear and cytoplasmic fractions. Protein samples were simultaneously obtained from the identical cell lysates used to isolate and sequence mRNA. W: whole cell lysate; C: cytoplasmic fraction; N: nuclear fraction. (**c**) Lethal fraction measured 32 hours after exposure to 10 µM ⍺-amanitin in Pol II switchover cells. Mean ± SD shown, n = 5 independent biological replicates. (**d**) RNAseq from cytoplasmic and nuclear fractions re-combined to recapitulate the whole cell mRNA extract, validating the quality of the fractionation. (**e**) Comparison of drug-induced expression changes of cytoplasmic mRNA in U2OS (x-axis) and U2OS-PTBP1-KO (y-axis) cells, following 8-hour exposure to 1 µM triptolide. Dashed lines denote 2.5 standard deviations from the identity line (x = y), off-diagonals are highlighted, and Pearson correlation coefficient of all genes is shown. (**f**) As in (e), but specifically for nuclear mRNAs. (**g**) Mapping of outliers from (e) onto the chemo-genetic profile for triptolide. (**h**) As in (g) but for panel (f).

**Extended Data FSSig. 20:**
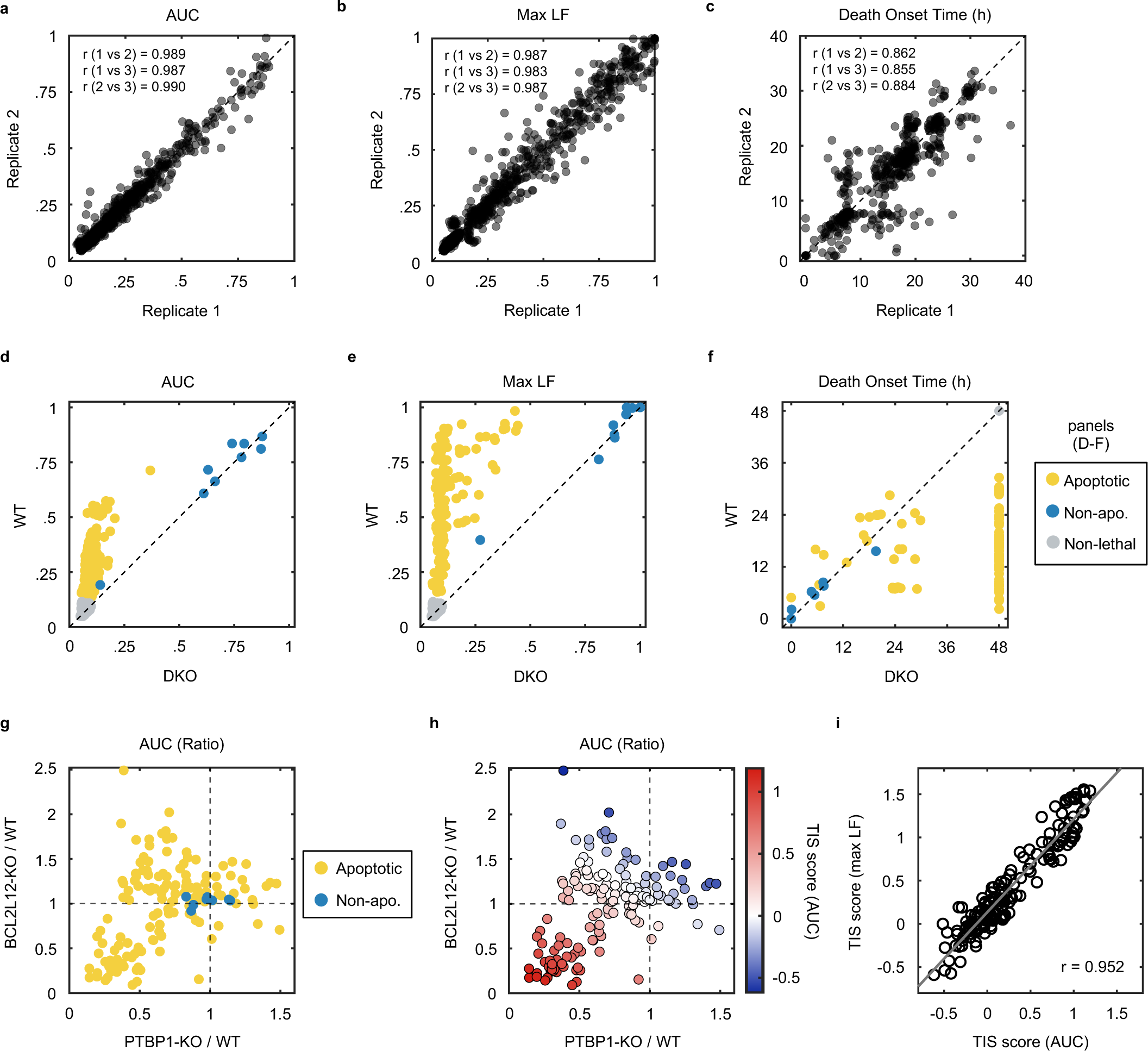
Evaluation of drug mechanism of killing using temporally resolved death kinetics. Related to. **Fig. 5**. (**a**) Quality control for the drug screen shown in Fig. 5. Correlation of AUC values between replicates. Each point represents a single drug-dose-genotype combination, measured using FLICK. Dashed line, x = y. Pearson correlation coefficient for each pair of replicates is shown. (**b**) As in (a), but for the maximum lethal fraction observed at assay endpoint (48 hours). (**c**) As in (a), but for the onset time of death. (**d-f**) Comparison of responses between *BAX/BAK1* DKO cells and U2OS cells for each drug-dose pair. Each condition is classified as apoptotic, non-apoptotic, or non-lethal. (**g**) Comparison of the change in cell death response following knockout of PTBP1 (x-axis) or BCL2L12 (y-axis) for each lethal drug-dose pair. Values of 1 denote no difference from parental U2OS cells. (**h**) Plot depicting how TIS scores vary across the response space. (**i**) AUC is a non-biased metric for generating TIS scores. TIS scores were calculated using max LF in an identical manner as AUC, producing similar results. Linear regression line is shown, along with the Pearson correlation coefficient.

**Extended Data FSSig. 21:**
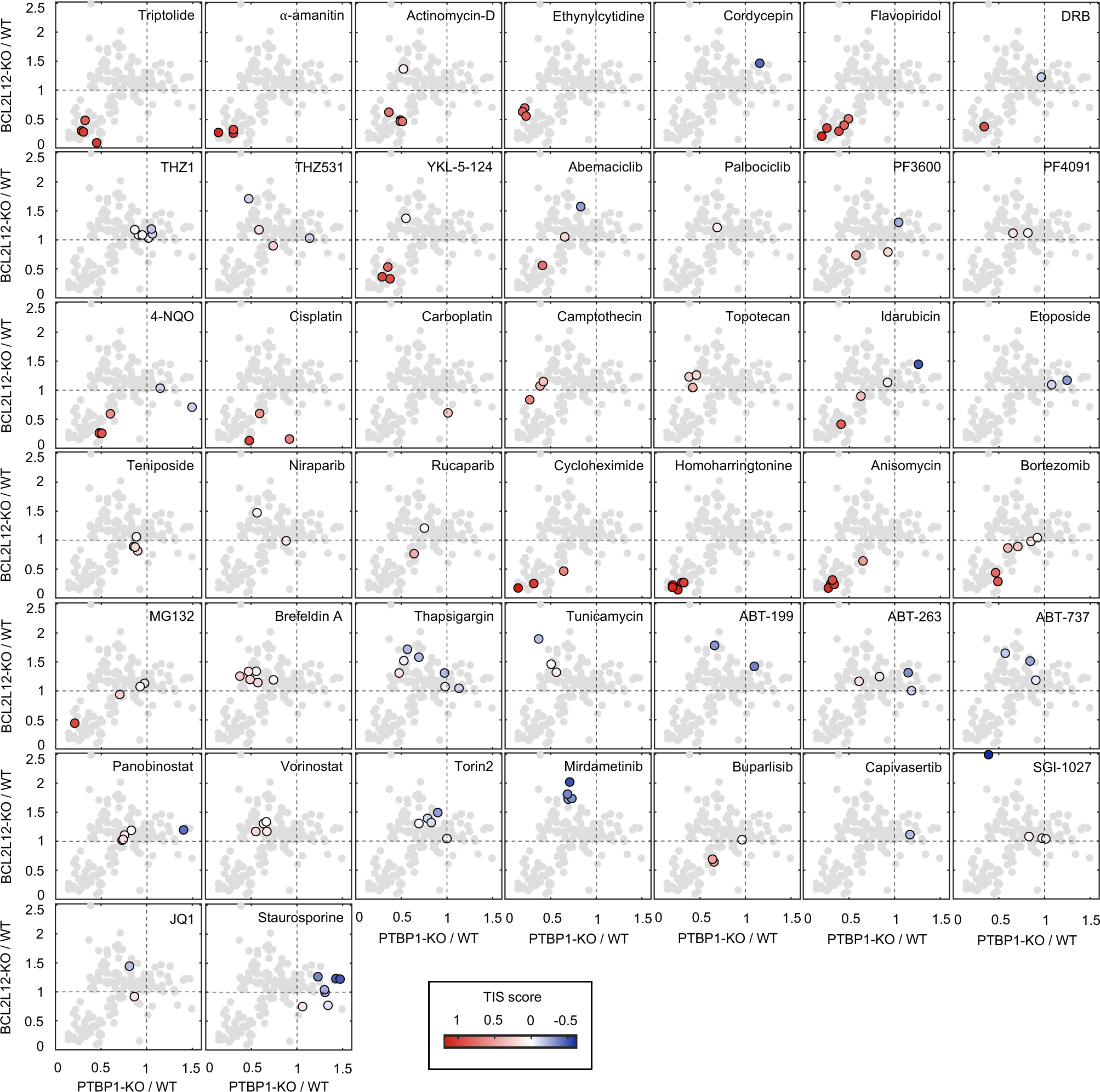
Evaluation of drug response using TIS score. Related to. **Fig. 5**. Map of the functional effect of PTBP1 and BCL2L12 knockout on the cell killing of each drug-dose pair (gray points). For each lethal drug, lethal doses are shown and colored by TIS score. Color bar applies to all panels.

**Extended Data FSSig. 22:**
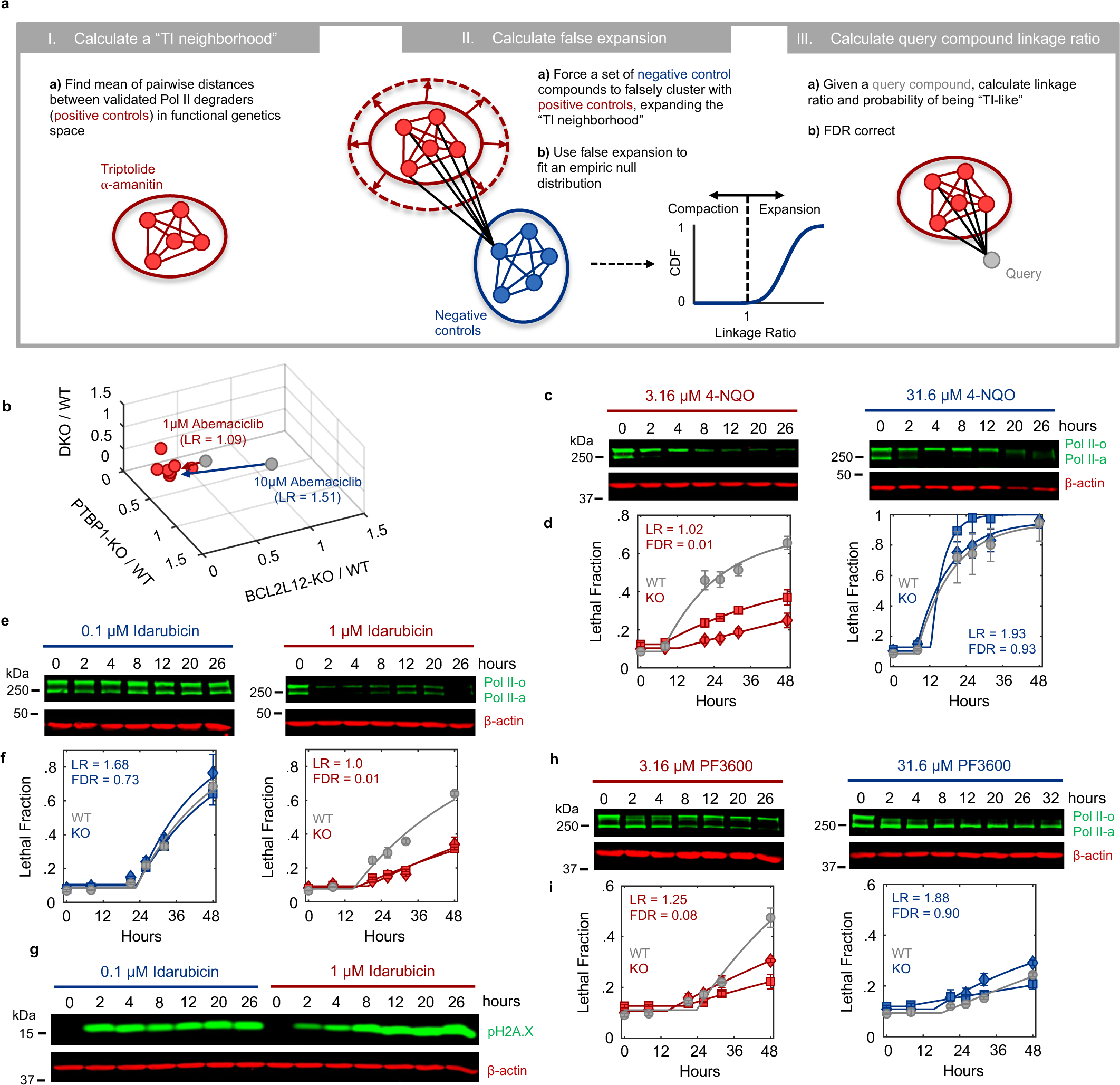
A statistical classifier to identify “TI-like” compounds. Related to. **Fig. 5**. (**a**) Conceptual overview of the probabilistic nearest neighbors-based classification approach (adapted from Pritchard, et al., 2013). (**b**) Classification of Abemaciclib. Triptolide and ⍺-amanitin (positive controls) are shown in red, Abemaciclib (query drug) is shown in gray. 1 µM Abemaciclib is classified as “TI-like”, whereas the 10 µM dose is not. Linkage ratios (LR) are shown. (**c**) Immunoblots of Rpb1 protein levels in U2OS cells following exposure to two different doses of 4NQO (3.16 µM left, 31.6 µM right). (**d**) Lethal fraction kinetics for 1 µM Abemaciclib (left) and 10 µM Abemaciclib (right) in U2OS cells (gray circles), PTBP1-KO cells (squares) and BCL2L12-KO cells (diamonds), measured using FLICK. Linkage ratios and associated FDR values are shown. Data are associated with blots above (respectively) in (C). (**e-f**) As in (c-d) for 0.1 µM and 1 µM Idarubicin. (**g**) Similar DDR signaling, as measured by pH2A.X levels in immunoblot, are seen at both doses of Idarubicin shown in (e-f). (**h-i**) As in (c-d) for 3.16 µM and 31.6 µM PF3600. For all panels TI-like drugs are red, Not TI-like are blue. For all panels with error bars, data are mean ± SD, n = 3 independent biological replicates.

**Extended Data FSSig. 23:**
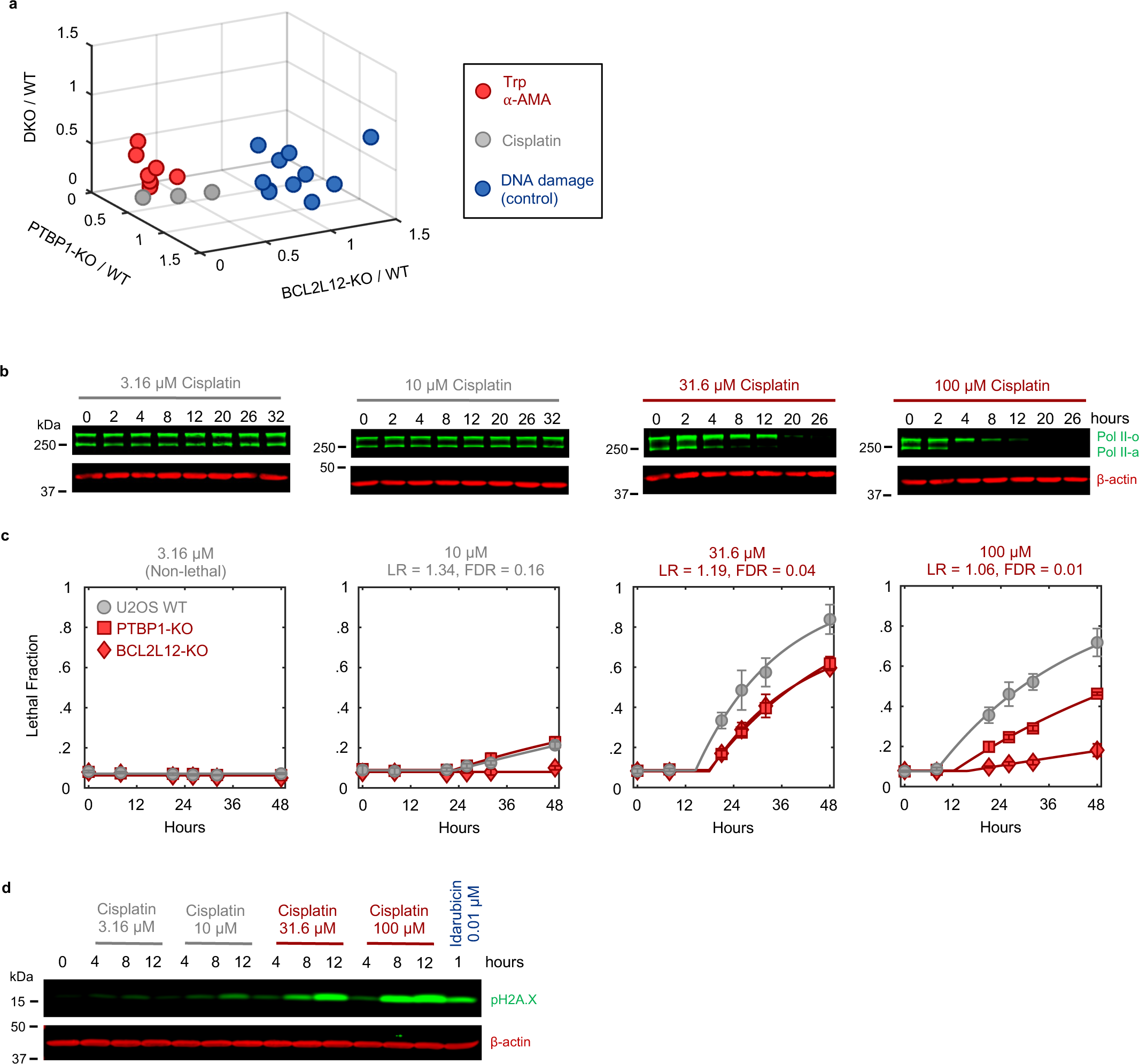
Lethal doses of Cisplatin require PTBP1 and BCL2L12 for cell killing and degrade Pol II protein. Related to. **Fig. 5**. (**a**) Classification of Cisplatin. Triptolide and ⍺-amanitin (positive controls) are shown in red, Cisplatin (query drug) is shown in gray. Other DNA damaging agents are shown in blue (Etoposide, Teniposide, Topotecan, and low-dose Idarubicin). (**b**) Immunoblots depicting Rpb1 protein levels over time following exposure to a range of Cisplatin doses. Images are representative of 3 independent biological replicates. (**c**) Lethal fraction kinetics of Cisplatin in U2OS, PTBP1-KO, and BCL2L12-KO cells, assessed at doses from (b). Linkage ratios and associated FDR values are shown. Data were collected in FLICK, and data are mean ± SD, n = 3 independent biological replicates. (**d**) Immunoblot of p-H2A.X levels across doses of Cisplatin. All doses of Cisplatin that induce appreciable amounts of p-H2A.X signaling are triptolide-like. A low, and non-lethal, dose of the DNA damaging agent Idarubicin is shown for comparison.

**Extended Data FSSig. 24:**
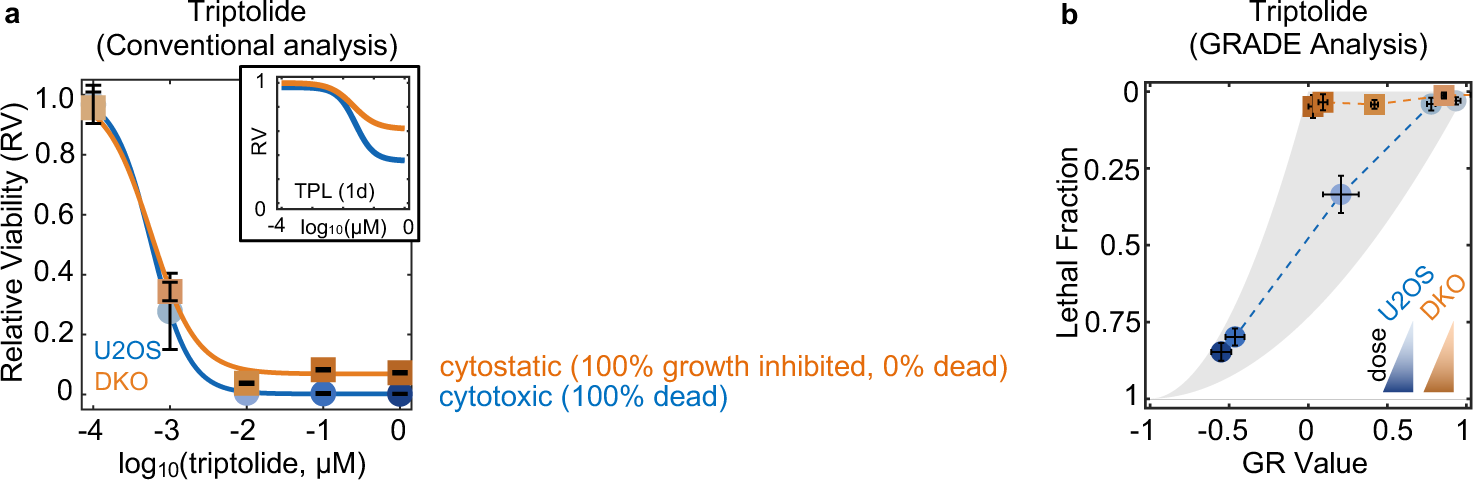
Conventional analysis metrics are insensitive to cell death and mask the existence of PDAR. (**a**) Triptolide (TPL) sensitivity in U2OS and DKO cells quantified using the conventional measure of drug sensitivity, Relative Viability (RV). RV was measured using the STACK assay 120 hours after drugging, or after 24 hours (inset). (**b**), GRADE analysis to infer the drug-induced growth rate and rate death rate from measurements of the lethal fraction and net population growth rate (GR value). Data for U2OS and DKO cells treated with triptolide for 48 hours, measured using STACK. For all panels with error bars, data are mean ± SD, n = 3 independent biological replicates.

